# Deep Annotation of Protein Function across Diverse Bacteria from Mutant Phenotypes

**DOI:** 10.1101/072470

**Authors:** Morgan N. Price, Kelly M. Wetmore, R. Jordan Waters, Mark Callaghan, Jayashree Ray, Jennifer V. Kuehl, Ryan A. Melnyk, Jacob S. Lamson, Yumi Suh, Zuelma Esquivel, Harini Sadeeshkumar, Romy Chakraborty, Benjamin E. Rubin, James Bristow, Matthew J. Blow, Adam P. Arkin, Adam M. Deutschbauer

**Author notes:** To whom correspondence should be addressed: MJB APA AMD. Website for interactive analysis of mutant fitness data: http://fit.genomics.lbl.gov/. Website with supplementary information and bulk data downloads: http://genomics.lbl.gov/supplemental/bigfit/.

## Abstract

**Summary:** The function of nearly half of all protein-coding genes identified in bacterial genomes remains unknown. To systematically explore the functions of these proteins, we generated saturated transposon mutant libraries from 25 diverse bacteria and we assayed mutant phenotypes across hundreds of distinct conditions. From 3,903 genome-wide mutant fitness assays, we obtained 14.9 million gene phenotype measurements and we identified a mutant phenotype for 8,487 proteins with previously unknown functions. The majority of these hypothetical proteins (57%) had phenotypes that were either specific to a few conditions or were similar to that of another gene, thus enabling us to make informed predictions of protein function. For 1,914 of these hypothetical proteins, the functional associations are conserved across related proteins from different bacteria, which confirms that these associations are genuine. This comprehensive catalogue of experimentally-annotated protein functions also enables the targeted exploration of specific biological processes. For example, sensitivity to a DNA-damaging agent revealed 28 known families of DNA repair proteins and 11 putative novel families. Across all sequenced bacteria, 14% of proteins that lack detailed annotations have an ortholog with a functional association in our data set. Our study demonstrates the utility and scalability of high-throughput genetics for large-scale annotation of bacterial proteins and provides a vast compendium of experimentally-determined protein functions across diverse bacteria.

## Background

Tens of thousands of bacterial genomes have been sequenced, revealing the predicted amino acid sequences of millions of distinct proteins. In sharp contrast, only a small proportion of these proteins have been studied experimentally and for most, presumptive functions can only be predicted via their similarity to experimentally characterized proteins. However, about one third of bacterial proteins are not similar enough to any characterized protein to be annotated by this approach. Furthermore, these predictions are often incorrect as homologous proteins may have different substrate specificities ^1^. This sequence-to-function gap represents a growing challenge for microbiology, because new bacterial genomes are being sequenced at an ever-increasing rate, while experimental protein characterization continues to be relatively slow ^2^.

One approach for determining an unknown protein’s function is to assess the consequences of a loss-of-function mutation of the corresponding gene under a large number of conditions ^3-6^. Transposon mutagenesis followed by sequencing (TnSeq) makes it possible to apply this principle more systematically and to identify mutant phenotypes at a genome-wide scale. In this approach, a transposon is used to create a complex library of mutant strains, each with a different random insertion site in the genome. If an insertion lies within a protein-coding gene, it disrupts the respective protein’s function. DNA sequencing can identify where each transposon has inserted and quantify the abundance of each mutant strain. Importantly, pools of hundreds of thousands of different mutants can be grown together, enabling genome-wide measurements of phenotypes and protein function in a single experiment ^7,8^. Coupling the approach to random DNA barcoding (RB-TnSeq) of individual mutants further increases the efficiency, facilitating the evaluation of a given mutant library under a large number of experimental conditions ^9^. Here, we use RB-TnSeq to directly address the sequence-to-function gap by systematically exploring the phenotypes of thousands of proteins from 25 bacteria under hundreds of experimental conditions (**Fig. 1a**).

**Figure 1.**
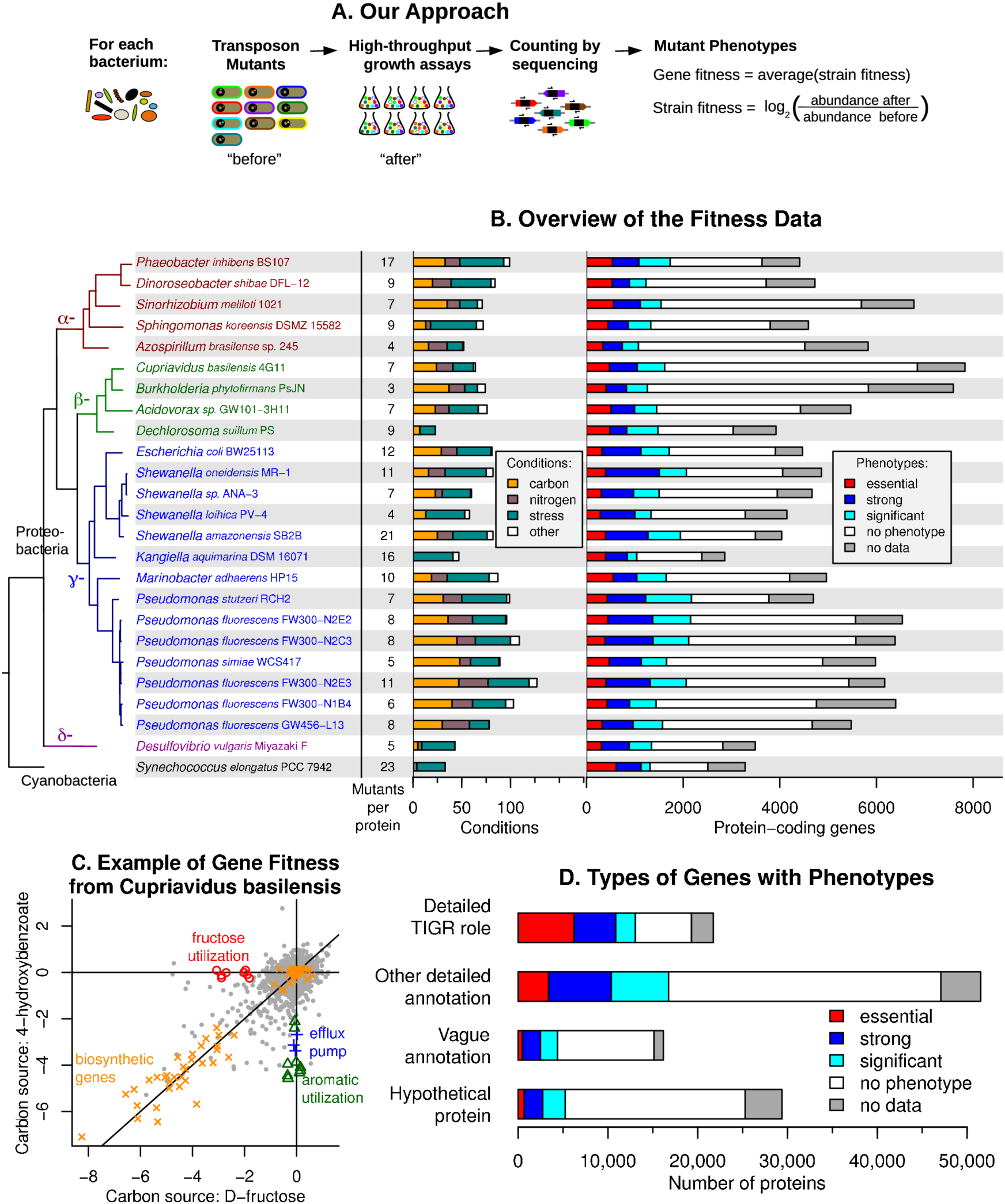
High-throughput genetics for 25 bacteria. (A) Our approach for measuring gene fitness. (B) Overview of our data. For each bacterium, we show the number of mutant strains that we used to estimate fitness for a typical protein (Methods), the types of conditions that we studied, and how many proteins had mutant phenotypes. A strong phenotype is defined as fitness < −2. (C) Gene fitness during the utilization of two carbon sources by *Cupriavidus basilensis*. See **Supplementary Table 5** for details on the highlighted genes. The 4-934 hydroxybenzoate data is the average from two biological replicates. (D) We classified the proteins from all bacteria by how informative their annotations were (Methods), and for each class, we show how many proteins have each type of phenotype.

## Results

### Mutant fitness compendia for 25 bacteria

To perform a systematic assessment of protein functions across a phylogenetically diverse set of bacteria, we chose 25 genetically tractable bacteria representing five different bacterial divisions and 16 different genera. In addition to the model bacterium *Escherichia coli* and 22 other aerobic heterotrophs, we studied a strictly anaerobic sulfate-reducing bacterium (*Desulfovibrio vulgaris*) and a strictly photosynthetic cyanobacterium (*Synechococcus elongatus*) (**Fig. 1b**). We generated a randomly barcoded transposon mutant library in each of the 25 bacteria, seven of which were previously described ^9-11^. For each mutant population, we used TnSeq to generate genome-wide maps of transposon insertion locations (Methods). Mutant populations were highly complex with 28,252 to 504,158 uniquely barcoded transposon insertions per genome, corresponding to 5 to 66 insertions for the typical (median) protein-coding gene. Genes that have very few or no transposon insertions are likely to be essential for viability in the conditions that were used to select the mutants. We identified 289 to 614 essential proteins per bacterium (6-23% of the genes in each), with 10,766 essential proteins total (Methods; **Supplementary Table 1; Supplementary Note 1**). To assess how many of these essential proteins are poorly annotated, we classified all existing protein annotations for each genome as “Detailed TIGR role” (for subfamilies that are annotated with functional roles by TIGRFAMs), “Other detailed” (for other functional annotations), “Vague” (for annotations like “transporter”), or “Hypothetical” (for functionally uninformative annotations) (Methods, **Supplementary Fig. 1**). Across the 25 bacteria, we identified 667 hypothetical proteins and 480 proteins with vague annotations that were essential (**Supplementary Table 1**).

To identify conditions suitable for mutant fitness profiling, we tested the growth of the wild-type bacteria in a wide range of conditions, including the utilization of 94 different carbon sources and 45 different nitrogen sources, and their inhibition by 55 stress compounds including antibiotics, surfactants, and heavy metals (**Supplementary Tables 2-4**). If we identified utilization or inhibition, then we performed a genome-wide fitness experiment using the corresponding mutant library. In the typical experiment, we grew a pool of mutants for 4-8 generations and used DNA barcode sequencing (BarSeq) ^12^ to compare the abundance of the mutants before and after growth (**Fig. 1a**). For each gene, we defined gene fitness to be the log_2_ change in abundance of mutants in that gene during the experiment (Methods, **Fig. 1a**).

For each bacterium, we successfully assayed fitness in 27 to 129 different experimental conditions (**Fig. 1b**), with a total of 2,035 bacterium-condition combinations. These conditions included the growth conditions discussed above as well as growth under varying pH and temperature and motility on agar plates. Including replicates, we conducted a total of 3,903 genome-wide fitness experiments that met our criteria for biological and internal consistency ^9^, and we obtained 14.9 million gene fitness values for 96,024 different non-essential protein-coding genes. The conditions and the fitness data can be viewed at the Fitness Browser (http://fit.genomics.lbl.gov/). Taken together, our results provide high-resolution genome-wide fitness maps for all 25 bacteria examined.

To establish the consistency of our data with known protein functions, we examined fitness data for the three most common classes of experiments: carbon utilization, nitrogen utilization, and stress. Genes without phenotypes have fitness values near 0, genes that are important for fitness have fitness values less than 0, and genes that are detrimental to fitness have fitness values greater than 0 (**Fig. 1c**). For the utilization of D-fructose or 4-hydroxybenzoate as the sole source of carbon in *Cupriavidus basilensis*, the fitness data identifies expected proteins for the catabolism of each substrate (**Fig. 1c**). Similarly, key enzymes and transporters required for the utilization of D-alanine or cytosine as the sole nitrogen source in *Azospirillum brasilense* are identified (**Supplementary Fig. 2a**). Lastly, in *Shewanella loihica*, orthologs of the CzcCBA heavy metal efflux pump ^13^ and the zinc responsive regulator ZntR are important for fitness in the presence of elevated zinc (**Supplementary Fig. 2b**). In addition to identifying expected proteins, in all of the presented examples we also identified proteins that were previously not known to be involved in the respective process, including an efflux pump important for 4-hydroxybenzoate utilization by *C. basilensis* (**Fig. 1c**, **Supplementary Table 5**). In summary, these examples highlight how the fitness data can validate protein annotations and identify new proteins involved in diverse biological processes.

We identified a significant mutant phenotype (false discovery rate < 5%) in at least one condition for 28,674 of the 96,024 (30%) non-essential proteins that we collected data for (**Fig. 1b**). Proteins with high or moderate similarity to another protein in the same genome (paralogs, alignment score above 30% of the self-alignment score) are less likely to have a phenotype (25% vs. 31%, *P* < 10^−15^; **Supplementary Fig. 3**), which likely reflects genetic redundancy. 18% of all proteins with fitness measurements were detrimental to fitness in at least one condition (**Supplementary Fig. 3**), which is consistent with previous reports that many proteins are detrimental in some conditions ^3,14^. Proteins annotated with a detailed TIGR role were particularly likely to have phenotypes, with more than half (52%) of those with fitness data showing a significant phenotype (**Fig. 1d**). In contrast, proteins that are not annotated with a detailed function (vague or hypothetical annotations) were less likely to have phenotypes (27% or 19%, respectively). Nevertheless, our assays identified phenotypes for 4,585 hypothetical proteins and for 3,902 other proteins with vague annotations (**Fig. 1d**). These include 2,828 proteins that do not belong to any characterized family in either Pfam or TIGRFAMs ^15,16^. Overall, we identified mutant phenotypes for 8,487 proteins that are not currently annotated with a detailed function, highlighting that a substantial proportion of hitherto vaguely annotated or hypothetical proteins have critical functions in bacteria.

### Conserved phenotypes are accurate predictors of protein function

To illuminate the biological function of individual proteins using genome-wide fitness data from multiple bacteria across a large number of experimental conditions, we used two principal approaches: (1) Identification of “specific” phenotypes that are observed only under one or a small number of conditions; (2) “Cofitness” patterns, where multiple proteins show similar fitness profiles across all conditions. Furthermore, we identified conserved specific phenotypes and conserved cofitness by comparing the data from 25 bacteria; and we tested the reliability of these conserved associations for annotating protein function.

To assign specific phenotypes, we identified proteins that had a significant and notable phenotype (|fitness| > 1) under only one or very few conditions (Methods). For example, the fluoride efflux protein CrcB ^17^ is required for fitness under elevated fluoride stress in multiple bacteria, but not for fitness in any of the hundreds of other experimental conditions that we tested (**Fig. 2a**). Among all genes with a significant phenotype under any condition, 34% have a specific phenotype (9,905 proteins), while the remaining genes either have weak but significant phenotypes (|fitness| < 1) or complex phenotypic patterns.

**Figure 2.**
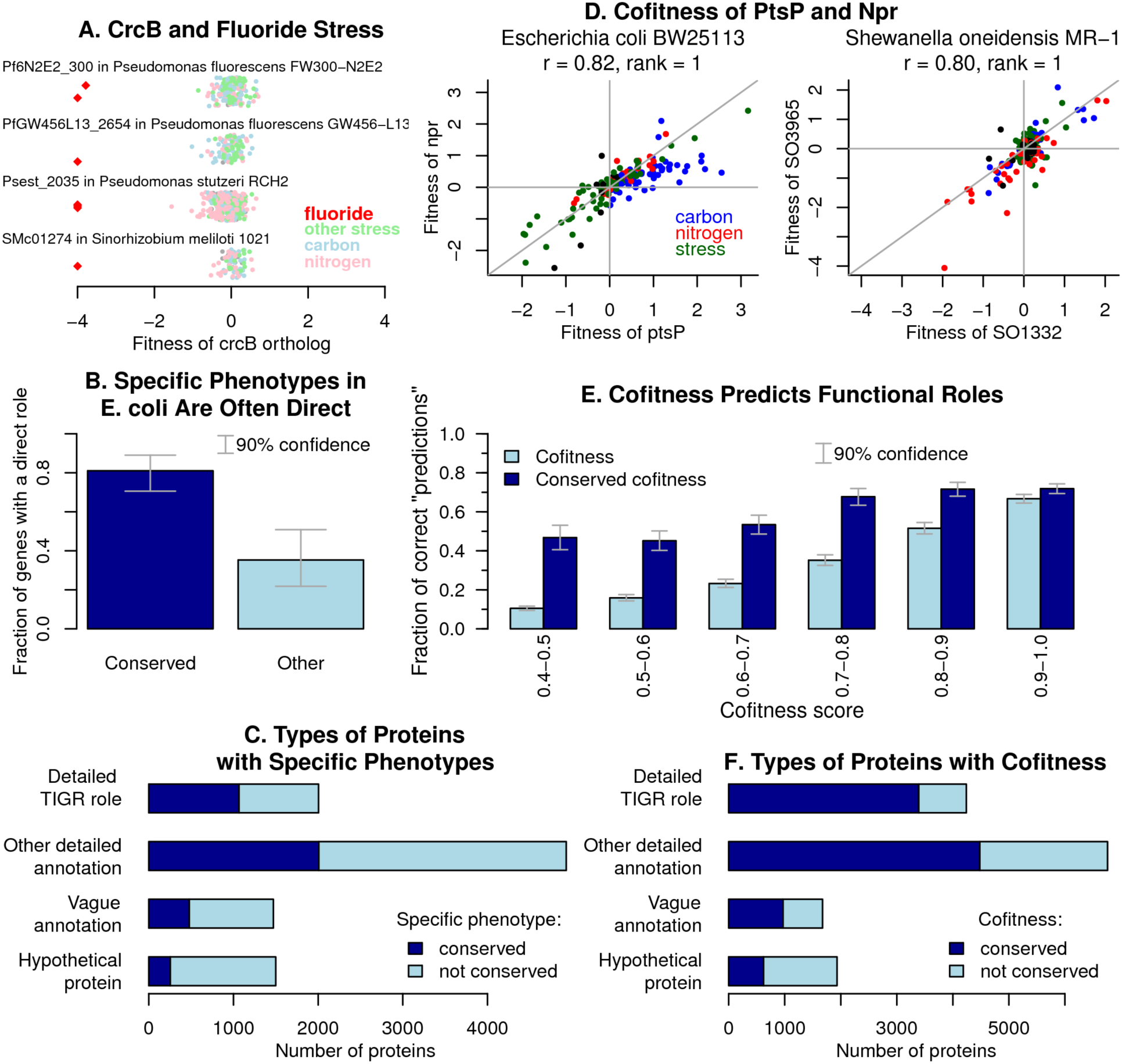
Identification of conserved phenotypes. (A) An example of a conserved and specific phenotype. Each point shows the fitness of *crcB* in an experiment, with fluoride stress experiments highlighted in red. Values less than −4 are shown at −4. The y-axis is random. (B) The fraction of *E. coli* proteins with a specific phenotype in defined media that are ‘directly’ involved in the uptake or catabolism of the compound. Conserved indicates that an ortholog of the *E. coli* protein from another bacterium is important for fitness during growth with the same compound (fitness < −1). (C) How many proteins of each type had a conserved specific phenotype or a specific phenotype. (D) Comparison of fitness values for *ptsP* and *npr* from *E. coli* and *S. oneidensis* across all tested conditions. The experiments are color coded by type. (E) Using TIGR subroles to test the accuracy of the gene-gene associations. (F) How many proteins of each type had at least one association from conserved cofitness (*r* > 0.6 in both bacteria) or else from cofitness (*r* > 0.8).

To assess whether specific phenotypes are useful indicators of protein function, we used our understanding of *E. coli* physiology and asked whether specific phenotypes in this bacterium could be used to accurately assign proteins to a “direct” enzymatic, regulatory, or transport function in a catabolic pathway. For example, all 4 proteins in *E. coli* with a specific phenotype during growth on D-xylose are directly involved in its utilization (*xylABFR*). More broadly, we manually examined 101 experimentally characterized proteins according to the EcoCyc database ^18^ with specific phenotypes on individual carbon and nitrogen sources and found that 67% have a known direct function in either the uptake or catabolism of that compound, or in the activation of genes that are (**Supplementary Table 6**). We also expected that if the specific phenotype were conserved (an orthologous protein from another bacterium has a notable phenotype in the same condition, see Methods), the association would be more reliable. Indeed, among the same 101 *E. coli* proteins, we found that 87% of proteins with conserved specific phenotypes were directly involved in utilization, as compared to 35% of proteins with specific phenotypes that were not conserved (**Fig. 2b**; *P* < 10^−4^, Fisher exact test; **Supplementary Table 6**). Thus, if a protein has a specific phenotype in a defined medium, it is likely to be directly involved in utilizing that substrate and hence a protein function can be assigned, and the probability is higher if the phenotype is conserved in another bacterium.

Overall, we identified specific phenotypes and conserved specific phenotypes for proteins of all annotation classes (**Fig. 2c**). In particular, specific phenotypes linked 2,970 proteins with vague or hypothetical annotations to over 100 diverse conditions, including 79 carbon sources, 42 nitrogen sources, and 54 stresses. These include conserved specific phenotypes, and hence high-confidence associations, for 733 such poorly annotated proteins.

Our second strategy for inferring a protein’s role was based on the observation that proteins with related functions often have similar fitness patterns across multiple conditions, which we term “cofitness” ^3,4^. For example, Npr and PtsP of the nitrogen phosphotransferase system (PTS) in *E. coli* exhibit high cofitness (**Fig. 2d**), consistent with the known role of PtsP in the phosphorylation and activation of Npr ^19^. Taking advantage of our comprehensive fitness dataset across 25 bacteria, we can now systematically address whether conserved cofitness is a stronger indicator of function than cofitness in one bacterium. For example, the orthologs of Npr and PtsP also have high cofitness in *S. oneidensis* (**Fig. 2d**).

We first identified all pairs of proteins with highly correlated phenotypes within an organism (cofit proteins), and the subset of these with orthologous protein pairs that are cofit across two or more organisms (conserved cofit proteins). To test how accurately cofitness or conserved cofitness links together functionally-related proteins, we next identified protein pairs with existing detailed functional annotations (a TIGR subrole from the TIGRFAMs database of protein families ^16^) and determined the frequency with which the subrole annotation of a protein is accurately “predicted” by that of the highest scoring cofit protein that is not nearby (see Methods for details). High-scoring cofitness in a single bacterium leads to predictions of TIGR subroles that are mostly correct, but the accuracy decays rapidly as the cofitness score decreases (**Fig. 2e**). In contrast, for conserved cofitness, the decay is much slower (**Fig. 2e**). Furthermore, conserved cofitness is significantly more accurate for a given number of predictions: for example, the top 2,000 predictions from cofitness (*r*> 0.81 for gene pairs from one bacterium) have 62% agreement, while the top 2,000 predictions from conserved cofitness (*r* > 0.56 for gene pairs from both bacteria) have 69% agreement (*P* < 10^−5^, Fisher exact test). Using thresholds of *r* > 0.8 for cofitness or *r* > 0.6 for conserved cofitness, we identified at least one association from cofitness or conserved cofitness for 15% of the genes with fitness data and for 45% of genes with significant phenotypes. We identified associations for all types of proteins (**Fig. 2f**), including for 1,934 hypothetical proteins and for 1,674 other vaguely-annotated proteins. When the associations link proteins that lack detailed annotations to proteins with TIGR subroles, the top-level roles are diverse, with the most common role (“cellular processes”) accounting for just 21% of them.

Combining the specific phenotypes and cofitness-derived functional associations, we identified a functional relationship for 19,962 proteins including 5,512 proteins with vague or hypothetical annotations. For 1,914 of these poorly-annotated proteins, we identified conserved associations, which are expected to be particularly robust predictors of protein function (**Supplementary Table 7**). These results demonstrate the utility of specific phenotypes and cofitness for elucidating the diverse roles of thousands of poorly-annotated proteins and highlight the advantages of collecting genome-wide fitness data for multiple bacteria.

### Genetic overviews of biological processes across diverse bacteria

Genome-wide mutant fitness profiling of phylogenetically diverse bacteria across a wide range of biological conditions provides a comprehensive genetic overview of each biological condition studied. To illustrate this, we examined proteins with conserved, specific, and important phenotypes (fitness < 0) during cisplatin stress (**Fig. 3a**). Cisplatin reacts with DNA to form crosslinks that block DNA replication, so we expected that DNA repair proteins would be important for growth in this condition ^3^. Indeed, of the 57 protein families that were specifically important for resisting cisplatin in more than one bacterium, 28 are known to be involved in DNA repair including the UvrABC nucleotide excision repair complex and the RecFOR homologous recombination pathway (**Fig. 3a**, **Supplementary Table 8**). In addition, 6 of the other characterized families that have conserved sensitivity to cisplatin are involved in cell division or chromosome segregation (**Fig. 3a**), which is relevant because DNA damage can inhibit DNA replication and lead to filamentous cells ^20^.

**Figure 3.**
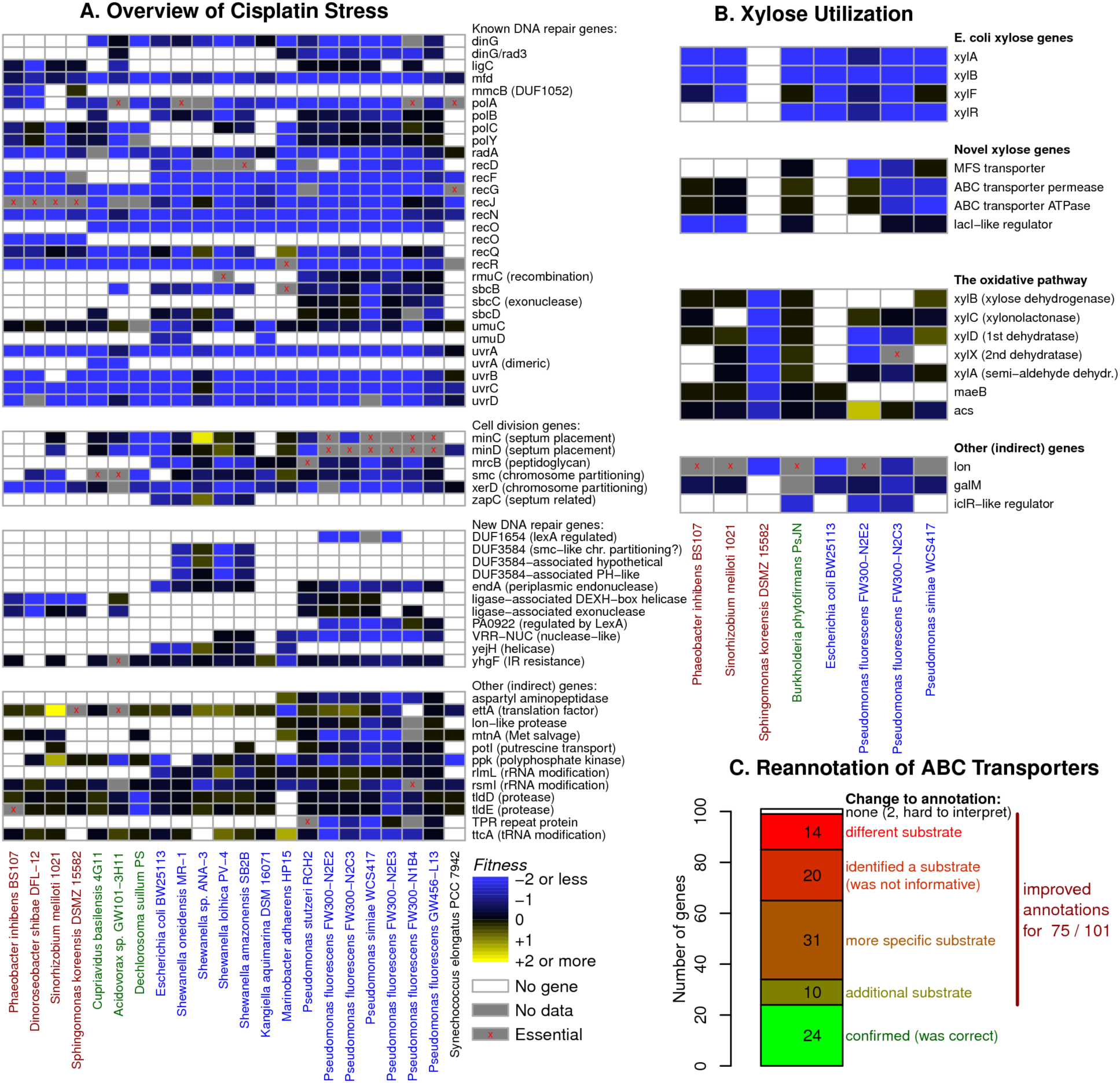
Genetic overviews for a condition or a class of proteins. (A) Overview of conserved specific phenotypes in cisplatin stress across bacteria. The data is the average for all successful cisplatin experiments for each bacterium, at up to 5 different concentrations. (B) Overview of specific phenotypes for the utilization of D-xylose as a carbon source in 8 bacteria. In the oxidative pathway, gene names for *xylABCDX* are from Stephens et al. ^27^ and should not be confused with *E. coli xylAB*, which are not related. In addition, putative orthologs are included in the heatmap even if they are not important for D-xylose utilization. For example, xylonolactonase is related to other lactonases that are important for the catabolism of other sugars. The data is the average of 1-2 replicate experiments for each bacterium. The color scale is the same as panel (A). (C) Summary of annotation improvements for ABC transporters based on an analysis of specific phenotypes.

We predict that 11 of the remaining 23 families with conserved cisplatin phenotypes are also involved in DNA repair, either because they contain DNA-related domains or because similar proteins are regulated by the DNA damage response regulator LexA in some bacteria ^21-23^. Three of the putative novel DNA repair genes (*endA, yejH, yhgF*) are present in the well-studied bacterium *E. coli,* and all 3 are important for resistance to ionizing radiation, which also damages DNA ^24^. This strongly suggests that their phenotypes are due to DNA damage. We also confirmed that an *E. coli* strain that has *endA* deleted is sensitive to cisplatin (**Supplementary Fig. 4**). The remaining 12 families likely have indirect roles in DNA repair, including 3 genes involved in tRNA or rRNA metabolism. Overall, our cisplatin experiments provide an overview of the proteins involved in DNA repair across diverse bacteria, including 28 protein families with a known role and 11 protein families whose putative role in DNA repair is not yet understood.

Similar genetic overviews were systematically obtained for the wide range of metabolic processes studied. To illustrate this, we identified at least one protein with a conserved, specific, and important phenotype in 67 carbon sources and 39 nitrogen sources. As an illustrative example, we examined D-xylose catabolism, which we assayed as the sole carbon source in 8 bacteria. We found that XylAB is required in *E. coli* and in 6 other bacteria, confirming its central and conserved role in D-xylose catabolism (**Fig. 3b**). In contrast, the well-characterized *E. coli* XylR regulator and XylF transporter are not required in each of the other 6 bacteria: two *Pseudomonads* use alternative transport proteins for D-xylose while *Phaeobacter inhibens* and *Sinorhizobium meliloti* require a lacI-like regulator for D-xylose utilization, as previously predicted ^9,25^. In contrast to the 7 bacteria that use the canonical XylAB pathway, we found that *Sphingomonas koreensis* uses an alternative, oxidative pathway for D-xylose utilization ^26,27^. Fitness data and comparative analysis to a similar pathway in archaea ^26,28^ suggest that *xylX* encodes the required enzyme 2-keto-3-deoxyxylonate dehydratase in *S. koreensis* (**Supplementary Table 9**). Our analysis also identified additional genes including *lon*, *galM*, and an *iclR*-like regulator that show conserved D-xylose utilization phenotypes across multiple bacteria, but whose exact roles in D-xylose catabolism remain to be elucidated. This comparative phenotypic analysis of D-xylose as a single carbon source represents just one of more than 100 carbon and nitrogen source experiments studied and highlights the power of our approach for the validation and discovery of new proteins and pathways involved in basic metabolic processes.

### Accurate annotations of individual protein families

Large-scale mutant fitness data can also be used to improve our understanding of proteins annotated with a general biochemical function but lacking substrate specificity. To illustrate this, we used the mutant fitness data to systematically reannotate the substrate specificities of 101 ABC transporter family proteins that have strong and specific phenotypes (fitness < −2 and statistically significant) during the utilization of diverse carbon or nitrogen sources (**Fig. 3c**; **Supplementary Table 10**). 20 of the 101 proteins only have vague annotations, and for these we made novel substrate predictions based on the specific phenotype data. For example, Dshi_0548 and Dshi_0549 from *Dinoroseobacter shibae* are annotated with no substrate specificity yet are important for utilizing xylitol. For another 31 proteins with moderately-specific substrate annotations, such as “amino acids”, we predicted a specific substrate within that group of compounds. For example, Ac3H11_2942 and Ac3H11_2943 from *Acidovorax* sp. 3H11 are annotated as transporting “various polyols”, whereas our data shows they are important for utilizing the polyol D-sorbitol but not the polyol D-mannitol. Another 14 proteins had incorrect annotations: for example, in three *Pseudomonads*, the L-carnitine transporter is mis-annotated as a choline transporter. In 10 cases, the specific phenotypes indicate that the protein transports a substrate that was not included in the annotation along with a substrate that was expected. For example, PS417_12705 from *P. simiae* is annotated as transporting D-mannitol, but this protein is also important for utilizing D-sorbitol and D-mannose. 24 proteins had correct annotations that were confirmed by our data; these included 22 cases in which the specific phenotype(s) were expected given the annotation, and 2 cases in which the gene has the expected mutant phenotype(s) but the association to a specific condition was misleading. The data for the remaining 2 proteins was hard to interpret. Overall, our fitness data provided improved annotations for 75 of the 101 transporters examined. This analysis also highlights how detailed computational annotations are often misleading: for 24 of the 50 ABC transporters that were annotated with a substrate (48%), the predicted specificity was erroneous or incomplete.

### Annotation of uncharacterized protein families

To evaluate the utility of fitness data for elucidating the functions of entirely uncharacterized protein families, we identified conserved cofitness or a conserved specific phenotype for 288 proteins that represent 77 different domains of unknown function (DUFs) from the Pfam database ^15^ (**Supplementary Table 11**). We manually examined the phenotypes of these 77 DUFs and we propose broad functional annotations for 13 of them and specific molecular functions for an additional 8 (**Supplementary Note 2**). For example, proteins containing UPF0126 are specifically important for glycine utilization in five bacteria (**Fig. 4a**). Since this family is predicted to be a membrane protein, we propose that it is more specifically a glycine transporter. As a second example, we found that members of the UPF0060 family of predicted transmembrane proteins have a conserved specific phenotype in the presence of elevated thallium in three bacteria (**Fig. 4b**). Consequently, we propose that UPF0060-containing proteins may function as a thallium-specific efflux pump. As a third example, we found that in three bacteria, DUF2849-containing proteins are cofit with an adjacent sulfite reductase (*cysI*) gene (**Fig. 4c**). The sulfite reductase is important for fitness in most defined media conditions because it is involved in sulfate assimilation, which was the only source of sulfur in our base media, and DUF2849 also seems to be involved in this process. These three bacteria lack *cysJ*, which is usually the electron source for *cysI*, and other bacterial genomes that contain DUF2849 also contain *cysI* but not *cysJ*. Based on this evidence, we propose that DUF2849 is an alternate electron source for sulfite reductase.

**Figure 4.**
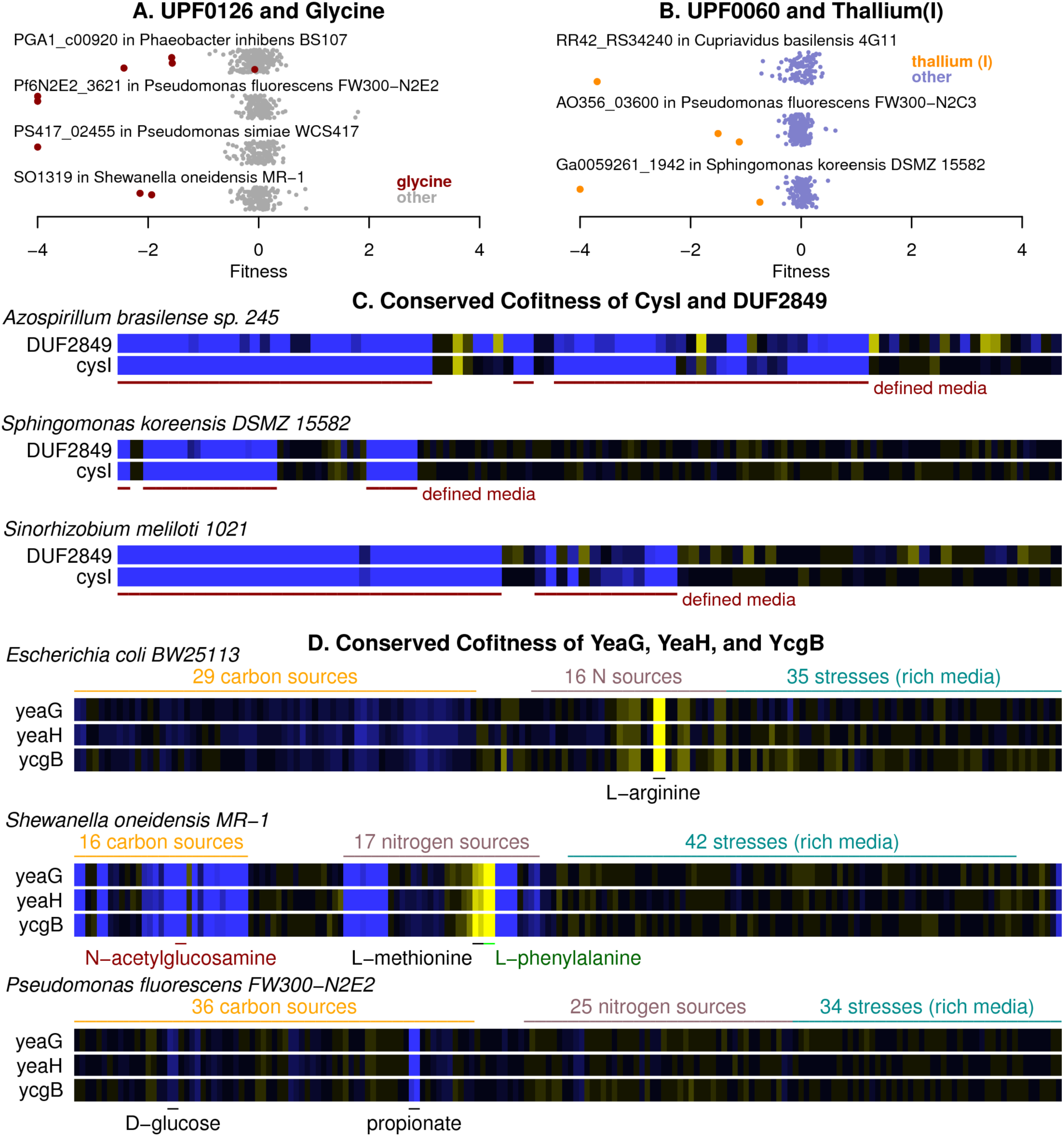
Annotation of proteins from uncharacterized families. (A, B) Conserved specific phenotypes for proteins of unknown function. Each point represents the fitness of the protein in an individual experiment. Values under −4 are shown at −4. The y-axis is random. (C) Heatmap of fitness data for *cysI* (sulfite reductase) and DUF2849 in three bacteria. (D) Heatmap of fitness data for *yeaG*, *yeaH*, and *ycgB* from three different bacteria. Certain experimental conditions with significant phenotypes are highlighted. The color scale is the same as in Fig. 3a.

Conserved phenotypic association can also be used to identify more complex pathways containing uncharacterized proteins. For example, the uncharacterized proteins YeaH and YcgB and the poorly characterized protein kinase YeaG ^29^ had high cofitness across six different bacteria (**Fig. 4d** shows the data from three of these bacteria; all *r > 0.7*). Interestingly, while these three proteins are more cofit with each other than with any other protein in each of the six bacteria, the phenotypes are not conserved across species (**Fig. 4d**). We validated some of these key phenotypes in growth assays with individual mutants. These experiments confirmed that in *E. coli*, mutants in all three genes have a growth advantage when L-arginine is the nitrogen source (**Supplementary Fig. 5**), whereas in *S. oneidensis*, mutants in all three genes grow slowly when N-acetylglucosamine is the carbon source (**Supplementary Fig. 6**). Based on these data and the protein kinase activity of YeaG, we propose that these three proteins act together in a conserved signaling pathway that is required for distinct cellular functions in different bacteria. Taken together, our data highlight how comprehensive fitness data can be used to provide novel experimental annotations for uncharacterized protein families and pathways.

### Relevance to all bacteria

Beyond the 25 bacteria experimentally studied in the present study, combining fitness data with comparative genomics offers the opportunity to assign phenotype-derived protein annotations across all sequenced bacterial genomes. To address this, we focused on the utility of our dataset to illuminate the functions of hypothetical or vaguely-annotated proteins from phylogenetically diverse bacterial genomes (**Methods**). For poorly-annotated proteins from these bacterial genomes, 14% have a putative ortholog (with at least 30% sequence similarity) with a significant phenotype in our dataset (**Fig. 5a**). Furthermore, for 11% of the poorly-annotated proteins, we can associate an ortholog (above 30% similarity) to a specific condition or to another protein with cofitness (**Fig. 5a**). Even at 30% similarity, our data should provide functional insight for many poorly characterized bacterial proteins. Supporting this, using gene ontology annotations supported by experimental evidence, Clark and Radivojac analyzed the similarity of molecular function between homologous proteins and found 60-70% functional conservation for homologs from anywhere from 30-100% identity ^30^. Not surprisingly, the probability of finding an ortholog with a functional association in our dataset is much higher for poorly-annotated proteins from bacterial divisions that we studied multiple representatives of (α,β,γ-Proteobacteria) than for such proteins from other bacteria (21% vs. 6%). For bacteria from these three divisions, we provide putative orthologs with functional associations for about 330 poorly-annotated proteins per average genome (3,913 proteins per genome * 40% poorly-annotated * 21%). To enable rapid access to the phenotypes and functional associations of the homologs of a protein of interest, we provide a “Fitness BLAST” web service (http://fit.genomics.lbl.gov/images/fitblast_example.html). The results from Fitness BLAST are available within the protein pages at IMG/M ^31^ and MicrobesOnline ^32^. We also provide a web page for comparing all of the proteins in a bacterium to the fitness data. Overall, our data provides functional information by homology for 14% of poorly annotated proteins across all sequenced bacteria.

**Figure 5.**
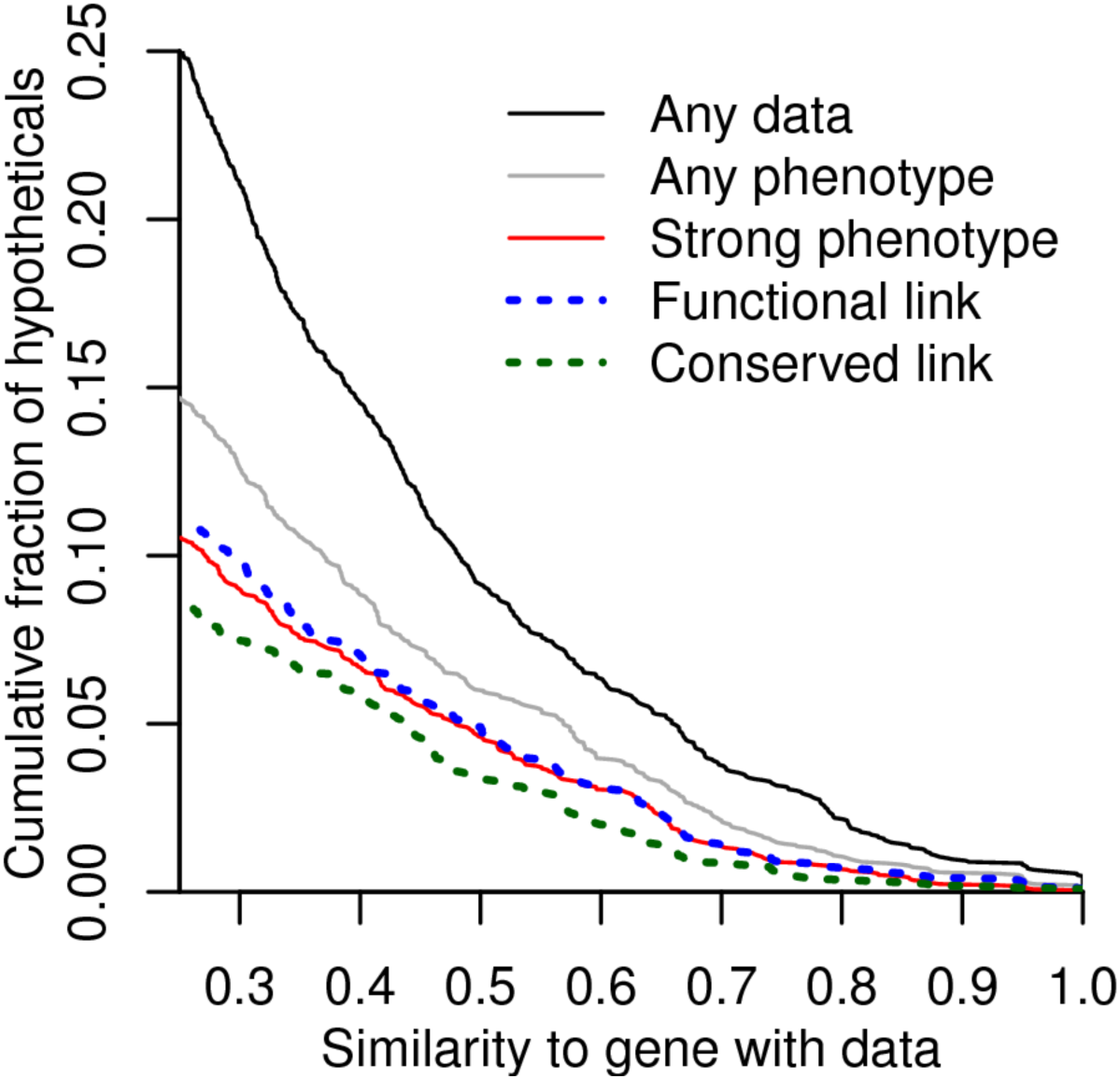
Relevance to all bacteria. We selected hypothetical or vaguely-annotated proteins from diverse bacterial species, we compared them to the genes that we have fitness data for (using protein BLAST), and we identified potential orthologs as best hits that were homologous over at least 75% of each protein’s length. We show the fraction of these proteins that have an ortholog with each type of phenotype and that is above a given level of sequence similarity. Similarity was defined as the ratio of the alignment’s bit score to the score from aligning the query to itself.

## Discussion

We have shown that high-throughput genetics can provide mutant phenotypes and functional associations for thousands of vaguely annotated and hypothetical bacterial proteins. Many of these functional associations were conserved, which increases the reliability of these associations. These associations can also highlight errors in current computational annotations, as we demonstrated for ABC transporters. Additionally, our comparative data provides unbiased functional overviews of biological processes by identifying proteins that are important for fitness under the same condition in multiple bacteria, as illustrated for D-xylose utilization and cisplatin stress.

The major challenge in extending our results to all bacterial proteins is their incredible diversity. Although we identified functional associations for 19,962 proteins and for 5,512 proteins that lack detailed annotations, these include putative orthologs of just 11% of the bacterial proteins that lack detailed annotations. Improving this coverage will require a larger effort to generate mutants in more diverse bacteria: our study included representatives of only 5 of the ∼40 divisions of bacteria that have been cultivated so far. Another challenge will be to improve the inference of protein function from phenotype. Many of the vaguely-annotated or incorrectly-annotated proteins belong to characterized families of enzymes or transporters and the main uncertainty as to their function is what substrate they act on. We proposed functional associations for 64% of the proteins that have significant phenotypes, and in principle, these associations could be used to automatically identify the physiological substrate for many of these proteins.

For thousands of proteins that previously lacked an informative annotation, our mutant phenotypes, and the functional associations derived from them, provide a rich resource to guide further study. To facilitate this, we developed the Fitness Browser web site (http://fit.genomics.lbl.gov) to view the fitness data for a gene or condition of interest. This site also supports the comparison of the fitness data across bacteria and incorporates tools for sequence-based annotation. In summary, our study demonstrates the scale with which large-scale fitness data can be collected in diverse bacteria and the utility of these data to provide insights into the functions of thousands of proteins, thereby helping to close the sequence-to-function gap in microbiology.

## Methods

### Bacteria

The bacteria mutagenized in this study are listed in **Supplementary Table 12**. Seven bacteria were isolated from groundwater collected from different monitoring wells at the Oak Ridge National Laboratory Field Research Center (FRC; http://www.esd.ornl.gov/orifrc/), and five have not been described previously: *Acidovorax* sp. GW101-3H11, *Pseudomonas fluorescens* FW300-N1B4, *P. fluorescens* FW300-N2E3, *P. fluorescens* FW300-N2C3, and *P. fluorescens* GW456-L13. *Acidovorax* sp. GW101-3H11 was isolated as a single colony on a Luria-Bertani (LB) agar plate grown at 30°C using an inoculum from FRC well GW101. *P. fluorescens* FW300-N1B4, *P. fluorescens* FW300-N2E3, and *P. fluorescens* FW300-N2C3 were all isolated at 30°C under anaerobic denitrifying conditions with acetate, propionate, and butyrate as the carbon source, respectively, using inoculum from FRC well FW300. *Pseudomonas fluorescens* GW456-L13 was isolated from FRC well FW456 under anaerobic incubations on a LB agar plate. We previously described the isolation of *Pseudomonas fluorescens* FW300-N2E2 ^33^ and *Cupriavidus basilensis* 4G11 ^34^. We also studied individual mutants of several organisms. For *E. coli* BW25113, we used single-gene deletions from the Keio collection ^35^. For *S. oneidensis* MR-1, we used transposon mutants that had been individually sequenced ^4^.

### Media and standard culturing conditions

A full list of the medias used in this study and their components are given in **Supplementary Table 13**. We routinely cultured *Acidovorax* sp. GW101-3H11, *Azospirillum brasilense* Sp245, *Burkholderia phytofirmans* PsJN, *Escherichia coli* BW25113, all *Pseudomonads* and *Shewanellae*, *Sinorhizobium meliloti* 1021, and *Sphingomonas koreensis* DSMZ 15582 in LB. *Cupriavidus basilensis* 4G11 was typically cultured in R2A media. *Dechlorosoma suillum* PS was cultured in ALP media ^11^. *Desulfovibrio vulgaris* Miyazaki F was grown anaerobically in lactate-sulfate (MOLS4) media, as previously described ^36,37^. We used marine broth (Difco 2216) for standard culturing of *Dinoroseobacter shibae* DFL-12, *Kangiella aquimarina* SW-154T, *Marinobacter adhaerens* HP15, and *Phaeobacter inhibens* BS107. *Synechococcus elongatus* PCC 7942 was normally cultured in BG-11 media with either 7,000 or 9,250 lux. All bacteria were typically cultured at 30°C except *Escherichia coli* BW25113 and *Shewanella amazonensis* SB2B, which were cultured at 37°C, and *P. inhibens* BS107, which was grown at 25°C. The *E. coli* conjugation strain WM3064 was cultured in LB at 37°C with diaminopimelic acid (DAP) added to a final concentration of 300 μM.

### High-throughput growth assays of wild-type bacteria

To assess the phenotypic capabilities of 23 aerobic heterotrophic bacteria and to identify conditions suitable for mutant fitness profiling, we monitored the growth of the wild-type bacterium in a 96-well microplate assay. These prescreen growth assays were performed in a Tecan microplate reader (either Sunrise or Infinite F200) with absorbance readings (OD_600_) every 15 minutes. All 96-well microplate growth assays contained 150 µL culture volume per well at a starting OD_600_ of 0.02. We used the grofit package in R ^38^ to analyze all growth curve data in this study. For carbon and nitrogen source utilization, we tested 94 and 45 possible substrates, respectively, in a defined medium (**Supplementary Tables 2 and 3**). We classified a bacterium as positive for usage of a particular substrate if (1) the maximum OD_600_ on the substrate was at least 1.5 greater than the average of the water controls and the integral under the curve (spline.integral) was 10% greater than the average of the water controls or (2) a successful genome-wide fitness assay was collected on the substrate, as described below. We included the second criterion because our automated scoring of the wild-type growth curves was conservative and did not include all conditions used for genome-wide fitness assays.

Additionally, for each of the 23 heterotrophic bacteria, we determined the inhibitory concentrations for 39-55 diverse stress compounds including antibiotics, biocides, metals, furans, aldehydes, and oxyanions. For each compound, we grew the wild-type bacterium across a 1,000-fold range of inhibitor concentrations in a rich media. We used the spline.integral parameter of grofit to fit dose-response curves and calculate the half-maximum inhibitory concentration (IC_50_) values for each compound (**Supplementary Table 4)**. For *Desulfovibrio vulgaris* Miyazaki F and *Synechococcus elongatus* PCC 7942, we did not perform these growth prescreen assays, rather, we just performed the mutant fitness assays across a broad range of inhibitor concentrations.

### Genome sequencing

We sequenced *Acidovorax sp.* GW101-3H11 and five *Pseudomonads* by using a combination of Illumina and Pacific Biosciences. For Illumina-first assembly, we used scythe (https://github.com/vsbuffalo/scythe) and sickle (https://github.com/najoshi/sickle) to trim and clean Illumina reads, we assembled with SPAdes 3.0 ^39^, we performed hybrid Illumina/PacBio assembly on SMRTportal using AHA ^40^, we used BridgeMapper on SMRTportal to fix misassembled contigs, we mapped Illumina reads to the new assembly with bowtie 2 ^41^, and we used pilon to correct local errors ^42^. For *Acidovorax sp.* GW101-3H11, we instead used A5 ^43^ to assemble the Illumina reads and we used AHA to join contigs together. For PacBio-first assembly, we used HGAP3 on SMRTportal, we used circlator to find additional joins ^44^, and we again used bowtie 2 and pilon to correct local errors. See **Supplementary Table 14** for a summary of these genome assemblies and their accession numbers. In addition, *Sphingomonas koreensis* DSMZ 15582 was sequenced for this project by the Joint Genome Institute, using Pacific Biosciences.

### Constructing pools of randomly barcoded transposon mutants

The transposon mutant libraries for seven bacteria were described previously ^9-11^. The other 18 bacteria were mutagenized with randomly barcoded plasmids containing a *mariner* or *Tn*5 transposon, a *pir-* dependent conditional origin of replication, and a kanamycin resistance marker, using the vectors that we described previously ^9^. The plasmids were delivered by conjugation with *E. coli* WM3064, which is a diaminopimelate auxotroph and is *pir*^+^. The conditions for mutagenizing each organism are described in **Supplementary Table 15**. Generally, we conjugated mid-log phase grown WM3064 donor (either *mariner* donor plasmid library APA752 or *Tn*5 donor plasmid library APA766) and recipient cells on 0.45 µM nitrocellulose filters (Millipore) overlaid on rich media agar plates supplemented with DAP. We used the rich medium preferred by the recipient (**Supplementary Table 15**). After conjugation, filters were resuspended in recipient rich media and plated on recipient rich media agar plate supplemented with kanamycin. After growth, we scraped together kanamycin resistant colonies into recipient rich media with kanamycin, diluted the culture back to a starting OD_600_ of 0.2 in 50-100 mL of recipient rich media with kanamycin, and grew the mutant library to a final OD_600_ of between 1.0 and 2.0. We added glycerol to a final volume of 10%, made multiple 1 or 2 mL −80°C freezer stocks, and collected cell pellets to extract genomic DNA for TnSeq. For *Desulfovibrio vulgaris* Miyazaki F, we selected for G418-resistant transposon mutants in liquid media with no plating step (**Supplementary Table 15**).

### Transposon insertion site sequencing (TnSeq)

Given a pool of mutants, we performed TnSeq just once to amplify and sequence the transposon junction and to link the barcodes to a location in the genome ^9^. We considered a barcode to be confidently mapped to a location if this mapping was supported by at least 10 reads. (For *Shewanella* sp. ANA-3, the threshold was 8 reads.) The number of unique barcodes (strains) mapped in each mutant library is shown in **Supplementary Table 15**. Given this mapping, the abundance of the strains in each sample can be determined by a simpler and cheaper protocol: amplifying the barcodes with PCR followed by barcode sequencing ^12^.

### Identifying essential genes

Genes that lack insertions or that have very low coverage in the start samples are likely to be essential or to be important for growth in rich media, as except for *Synechococcus elongatus*, pools of mutants were produced and recovered in media that contained yeast extract. We used previously published heuristics ^10^ to distinguish likely-essential genes from genes that are too short or that are too repetitive to map insertions in. Briefly, for each protein-coding gene, we computed the total read density in TnSeq (reads / nucleotides across the entire gene) and the density of insertion sites within the central 10-90% of each gene (sites/nucleotides). We then excluded genes that might be difficult to map insertions within because they were very similar to other parts of the genome (BLAT score above 50) and also very-short genes of less than 100 nucleotides. Given the median insertion density and the median length of the remaining genes, we asked how short a gene could be and still be unlikely to have no insertions at all by chance (*P* < 0.02, Poisson distribution). Genes shorter than this threshold were excluded; the threshold varied from 100 nucleotides for *Phaeobacter inhibens* to 600 nucleotides for *Desulfovibrio vulgaris* Miyazaki. For the remaining genes, we normalized the read density by GC content by dividing by the running median of read density over a window of 201 genes (sorted by GC content). We normalized the insertion density so that the median gene's value was 1. Protein-coding genes were considered essential or important for growth if we did not estimate fitness values for the gene and both the normalized insertion density and the normalized read density were under 0.2. A validation of this approach is described in **Supplementary Note 1**.

### Mutant fitness assays

For each mutant library, we performed competitive mutant fitness assays under a large number of growth conditions that were chosen based on the results of high-throughput growth assays of wild-type bacteria (see above). The full list of experiments performed for each mutant library is available at http://fit.genomics.lbl.gov. Our analysis includes 385 successful experiments from Wetmore *et al.* ^9^ and 36 successful experiments from Melnyk *et al.* ^11^. The other 3,482 successful fitness assays are described here for the first time. In general, all growth assays with carbon sources, nitrogen sources, and inhibitors were done as previously described ^9^. Briefly, an aliquot of the mutant library was thawed and inoculated into 25 mL of rich media with kanamycin and grown to mid-log phase in a flask. Depending on the mutant library, this growth recovery took between 3 and 24 hours. After recovery, we collected pellets for genomic DNA extraction and barcode sequencing (BarSeq) of the input or “start” sample. We used the remaining cells to set up multiple mutant fitness assays with diverse carbon and nitrogen sources in defined media and diverse inhibitors in rich media, all at a starting OD_600_ of 0.02. In addition, for most bacteria, we profiled growth of the mutant library at different pH and at different temperatures. After the mutant library grew to saturation under the selective growth condition (typically 4 to 8 population doublings), we collected a cell pellet for genomic DNA extraction and BarSeq of the “end” sample. As described below, we calculate gene fitness from the barcode counts of the end sample relative to the start sample.

We used a number of different growth formats and media formations for mutant fitness assays across the 25 bacteria. The complete metadata for all mutant fitness assays in each bacterium are available at the supporting website. A full list of compound components for each growth media are contained in **Supplementary Table 13**. Many fitness assays were done in 48-well microplates (Greiner) with 700 µL culture volume per well and grown in a Tecan Infinite F200 plate reader with OD_600_ measurements every 15 minutes. For these 48-well microplate assays, we combined the cultures from two replicate wells before genomic DNA extraction (total volume of experiment = 1.4 mL). For 24-well microplate experiments, we used deep-well plates with 1.5 mL (for inhibitors) or 2 mL (for carbon and nitrogen sources) total culture volume per well. All 24-well microplate experiments were grown in a Multitron incubating shaker (Innova). For 24-well microplate experiments, we typically took the OD_600_ of each culture every 12 to 24 hours in a Tecan plate reader (after transferring the cells to a Greiner 96-well microplate). Over 1,000 experiments, primarily carbon source and temperature experiments, were done in glass test tubes with 5 mL culture volumes. For the test tube experiments, we monitored OD_600_ every 12 to 24 hours with a standard spectrophotometer and cuvettes.

For stress experiments, we tried to use a concentration of each compound that allows growth (because if there is no growth, then the abundance of the strains will not change) but significantly inhibits growth (or else the fitness pattern is likely to be as if the compound were not added). Ideally, the concentration is such that the growth rate is cut in half. For aerobic heterotrophs, we measured growth across several orders of magnitude of concentrations of each stress compound, as described above. This gave a rough estimate of what concentration to use. Then, when we performed the fitness assays, we used a few different concentrations to try and capture an inhibitory but sub-lethal concentration. For assays done in 48-well microplates and grown in a Tecan Infinite F200 plate reader, we could confirm that the culture was inhibited relative to a no stress control. For stress assays in 24-well, deep-well microplates and grown in the Multitron shaker, we took OD readings approximately every 12 hours to estimate which cultures were inhibited. In practice, we often did multiple mutant fitness assays with different concentrations of the same inhibitor. We also collected fitness data in plain rich media without an added inhibitory compound.

For some carbon source experiments in *Desulfovibrio vulgaris* Miyazaki F, which is strictly anaerobic, we grew the mutant pool in 18 × 150 mm hungate tubes with a butyl rubber stopper and an aluminum crimp seal (Chemglass Life Sciences, Vineland, NJ) with a culture volume of 10 mL and a headspace of about 15 mL. For the remainder of the *Desulfovibrio vulgaris* fitness experiments, we grew the mutant pool in 24 well microplates inside of the anaerobic chamber. We used OD_600_ measurements to determine which cultures were inhibited by varying concentrations of stress compounds. Similarly, for six of the other heterotrophs, we measured gene fitness during anaerobic growth. All anaerobic media was prepared within a Coy anaerobic chamber with an atmosphere of about 2% H_2_, 5% CO_2_, and 93% N_2_.

For *Synechococcus elongatus* PCC 7942, which is strictly photosynthetic, we recovered the library from the freezer in BG-11 media at a light level of 7000 lux and we conducted fitness assays at 9250 lux. We used OD_750_ to measure the growth of *S. elongatus*. Most *S. elongatus* mutant fitness assays were done in the wells of a 12-well microplate (Falcon) with a 5 mL culture volume.

In addition to growth assays in liquid media, we successfully studied motility in 12 bacteria using a soft agar assay. For motility assays, the mutant pool was inoculated into the center of a 0.3% agar rich media plate and “outer” samples with motile cells were removed with a razor after 24-48 hours. In many instances, we also removed an “inner” sample of cells from near the point of inoculation. Not all bacteria we assayed were motile in this soft agar assay and others were motile but did not give mutant fitness results that passed our quality metrics.

In four bacteria, we also assayed survival. In these assays, a mutant pool was subjected to a stressful condition (either extended stationary phase or a low temperature of 4°C) for a defined period; then, to determine which strains are still viable, they were recovered in rich media for a few generations. After recovery in rich media, the cells were harvested for genomic DNA extraction and BarSeq.

### Barcode sequencing (BarSeq)

Genomic DNA extraction and barcode PCR were performed as described previously ^9^. Most genomic DNA extractions were done in a 96-well format using a QIAcube HT liquid handling robot (QIAGEN). We used the 98°C BarSeq PCR protocol ^9^, which is less sensitive to high GC content. In general, we multiplexed 48 samples per lane of Illumina HiSeq. For *E. coli*, we sequenced 96 samples per lane instead.

### Computation of fitness values

Fitness data was analyzed as previously described ^9^. Briefly, the fitness value of each strain (an individual transposon mutant) is the normalized log_2_(strain barcode abundance at end of experiment/strain barcode abundance at start of experiment). The fitness value of each gene is the weighted average of the fitness of its strains; only strains that have sufficient start reads and lie within the central 10-90% of the gene are included. The median number of usable strains per gene in each bacterium is shown in **Fig. 1b.** The gene fitness values were then normalized to remove the effects of variation in genes’ copy number: the median for each scaffold is set to zero, and for large scaffolds, the running median of the gene fitness values is subtracted. Also, for large scaffolds, the peak of the distribution of gene fitness values is set to be at zero. For example, a gene fitness value of −2 means that the strains with transposon mutant insertions in that gene were, on average, at 25% of their original abundance at the end of the experiment.

Fitness experiments were deemed successful using the quality metrics that we described previously ^9^. These metrics ensure that the typical gene has sufficient coverage, that the fitness values of independent insertions in the same gene are consistent, and that there is no GC bias ^9^. Experiments that did not meet these thresholds were excluded from our analyses. The remaining experiments show good agreement between biological replicates, with a median correlation of 0.88 for gene fitness values from defined media experiments. Stress experiments sometimes have little biological signal, as they are usually done in rich media and mutants of genes that are important for growth in rich media may be absent from the pools. Nevertheless, the median correlation between replicate stress experiments was 0.68.

To estimate the reliability of the fitness value for a gene in a specific experiment, we use a *t*-like test statistic which is the gene’s fitness divided by the standard error ^9^. The standard error is the maximum of two estimates. The first estimate is based on the consistency of the fitness for the strains in that gene. The second estimate is based on the number of reads for the gene.

Even mild phenotypes were quite consistent between replicate experiments if they were statistically significant. For example, if a gene had a mild but significant phenotype in one replicate (0.5 < |fitness| < 2 and |*t*| > 4), then the sign of the fitness value was the same in the other replicate 95.5% of the time. Because this comparison might be biased if the two replicates were compared to the same control sample, only replicates with independent controls were included.

### Genes with statistically significant phenotypes

We averaged fitness values from replicate experiments. We combined *t* scores across replicate experiments with two different approaches. If the replicates did not share a start sample and were entirely independent, then we used *t*_comb_ = sum(*t*) / sqrt(*n*), where *n* is the number of replicates. But if the replicates used the same start sample then this metric would be biased. To correct for this, we assumed that the start and end samples have similar amounts of noise. This is conservative because we usually sequenced the start samples with more than one PCR and with different multiplexing tags. Given this assumption and given that variance(A+B) = variance(A) + variance(B) if A and B are independent random variables, it is easy to show that the above estimate of t_comb_ needs to be decreased by a factor of sqrt((*n*\*\*2 + *n*)/(2\**n*)), where *n* is the number of replicates.

Our standard threshold for a significant phenotype was |fitness| > 0.5 and |combined *t*| > 4, but this was increased for some bacteria to maintain a false discovery rate (FDR) of less than 5%. We use a minimum threshold on |fitness| as well as |*t*| to account for imperfect normalization or for other small biases in the fitness values. To estimate the number of false positives, we used control experiments, that is, comparisons between different measurements of different aliquots of the same start sample. However we did not use some previously-published control comparisons (from ^9^) that used the old PCR settings and had strong GC bias (to exclude these, we used the same thresholds that we used to remove biased experiments). The estimated number of false positive genes was then the number of control measurements that exceeded the thresholds, multiplied by the number of conditions and divided by the number of control experiments. As a second approach to estimate the number of false positives, we used the number of expected false positives if the *t_comb_* values follow the standard normal distribution (2 * P(*z* > *t*) * #experiments * #genes). If either estimate of the false discovery rate was above 5%, we raised our thresholds for both |fitness| and |*t_comb_*| in steps of 0.1 and 0.5, respectively, until FDR < 5%. The highest thresholds used were |fitness| > 0.9 and |t| > 6. Also, for *Pseudomonas fluorescens* m FW300-N1B4, we identified six genes with large differences between control samples (|fitness| ≈ 2 and |*t*| ≈ 6). These genes cluster on the chromosome in two groups and are strongly cofit, and several of the genes are annotated as being involved in capsular polysaccharide synthesis. Because this bacterium is rather sticky, we suspect that mutants in these genes are less adherent and were enriched in some control samples due to insufficient vortexing, so these six genes were excluded when estimating the number of false positives.

### Sequence analysis

To assign genes to Pfams ^15^ or TIGRFAMs ^16^, we used HMMer 3.1b1 ^45^ and the trusted score cutoff for each family. We used Pfam 28.0 and TIGRFAM 15.0. We used only the curated families in Pfam (“Pfam A”).

To identify putative orthologs between pairs of genomes, we used bidirectional best protein BLAST hits with at least 80% alignment coverage both ways. We did not use any cutoff on similarity, as a similarity of phenotype can show that distant homologs have conserved functions.

To measure the relevance of our fitness data to diverse bacteria, we started with 1,752 bacterial genomes from MicrobesOnline ^32^. We grouped together closely related genomes if they were separated from their common ancestor by less than 0.01 substitutions per site in highly conserved proteins (using the MicrobesOnline species tree). These groups correspond roughly to species. We selected one representative of each group at random, which gave us 1,236 divergent bacterial genomes. We selected 5 proteins at random, regardless of their annotation, from each of these genomes to form our sample of bacterial protein-coding genes. The best-scoring hit to one of the 25 bacteria that was studied was considered as a potential ortholog if the alignment coverage was 75% or more; we used a range of threshold for similarity but our recommended cutoff is that the ratio of the BLAST alignment score to the self score be above 0.3 (as in ^46^).

To estimate the evolutionary relationships of the bacteria that we studied (**Fig. 1b**), we used Amphora2 ^47^ to identify 31 highly-conserved proteins in each genome and to align them, we concatenated the 31 protein alignments, and we used FastTree 2.1.8 ^48^ to infer a tree.

### Specific phenotypes

We defined a specific phenotype for a gene in an experiment as: |fitness| > 1 and |*t*| > 5 in this experiment; |fitness| < 1 in at least 95% of experiments; and the fitness value in this experiment is noticeably more extreme than most of its other fitness values (|fitness| > 95th percentile(|fitness|) + 0.5).

We considered a specific phenotype to be conserved if a potential ortholog had a specific phenotype with the same sign in a similar experiment with the same carbon source, nitrogen source, or stressful compound (but not necessarily using the same base media or the same concentration of the compound). For specific-important phenotypes, we also considered a specific phenotype to be conserved if a potential ortholog had fitness < −1 and *t* < −4 in a similar experiment.

### “Predicting” TIGR subroles from cofitness or conserved cofitness

We only considered hits that were in the top 10 cofit hits for a gene, only hits with cofitness above 0.4 (or conserved cofitness above 0.4), and only hits that were at least 10 kilobases from the query gene. In these cases, we predict that the gene has the same subrole as the best-scoring hit. When testing cofitness, the hits were sorted by the cofitness. When testing conserved cofitness, the hits were sorted by the lower of the cofitness in this organism and the best cofitness for orthologs in other organisms.

### Genome and gene annotations

For previously-published genomes, gene annotations were taken from MicrobesOnline, Integrated Microbial Genomes (IMG), or RefSeq. Newly-sequenced genomes were annotated with RAST ^49^, except that *S. koreensis* DSMZ 15582 was annotated by IMG. See **Supplementary Table 16** for a summary of genome annotations and their accession numbers.

### Classification of how informative annotations are

To assess the existing computational annotations for these genomes, we classified all of their proteins into one of four groups: detailed TIGR role, hypothetical, vague, or other detailed. (1) “Detailed TIGR role” includes proteins that belong to a TIGRFAM role other than “Unclassified”, “Unknown function”, or “Hypothetical proteins” and had a subrole other than “Unknown substrate”, “Two-component systems”, “Role category not yet assigned”, “Other”, “General”, “Enzymes of unknown specificity”, “Domain”, or “Conserved”. (2) “Hypothetical” includes proteins containing the annotation description “hypothetical protein”, “unknown function”, “uncharacterized”, or if the entire description matched “TIGRnnnnn family protein” or “membrane protein”. (3) Proteins were considered to have “vague” annotations if the gene description contained “family”, “domain protein”, “related protein”, “transporter related”, or if the entire description matched common non-specific annotations (“abc transporter atp-binding protein”, “abc transporter permease”, “abc transporter substrate-binding protein”, “abc transporter”, “acetyltransferase”, “alpha/beta hydrolase”, “aminohydrolase”, “aminotransferase”, “atpase”, “dehydrogenase”, “dna-binding protein”, “fad-dependent oxidoreductase”, “gcn5-related n-acetyltransferase”, “histidine kinase”, “hydrolase” “lipoprotein”, “membrane protein”, “methyltransferase”, “mfs transporter”, “oxidoreductase”, “permease”, “porin”, “predicted dna-binding transcriptional regulator”, “predicted membrane protein”, “probable transmembrane protein”, “putative membrane protein”, “response regulator receiver protein”, “rnd transporter”, “sam-dependent methyltransferase”, “sensor histidine kinase”, “serine/threonine protein kinase”, “signal peptide protein”, “signal transduction histidine kinase”, “tonb-dependent receptor”, “transcriptional regulator”, “transcriptional regulators”, or “transporter”). The remaining proteins were considered to have “other detailed” annotations.

To identify a subset of the proteins annotated as “hypothetical” or “vague” that do not belong to any characterized families, we relied on Pfam and TIGRFAM. A Pfam was considered to be uncharacterized if its name began with either DUF or UPF (which is short for “uncharacterized protein family”). A TIGRFAM was considered to be uncharacterized if it had no link to a role or if the top-level role was “Unknown function”. To identify poorly-annotated proteins from diverse bacteria (for **Fig. 5a**), we used the rules for vague annotations only.

### Availability of data and code

See the Fitness Browser (http://fit.genomics.lbl.gov) or the data downloads page (http://morgannprice.org/bigfit/), which includes the supplementary information. The BarSeq or TnSeq reads were analyzed with the RB-TnSeq scripts (https://bitbucket.org/berkeleylab/feba); we used statistics versions 1.0.2, 1.0.3, or 1.1.0 of the code.

## Acknowledgements

We thank Axel Visel for help editing the manuscript and Victoria Lo, Wenjun Shao, and Keith Keller for technical assistance with the Fitness Browser web site. Sequencing was performed at: the Vincent J. Coates Genomics Sequencing Laboratory (University of California at Berkeley), supported by NIH S10 Instrumentation Grants S10RR029668, S10RR027303, and OD018174; at the DOE Joint Genome Institute; at the College of Biological Sciences ^UC^DNA Sequencing Facility (UC Davis); and at the Institute for Genomics Sciences (University of Maryland).

Studies of novel isolates were conducted by ENIGMA and were supported by the Office of Science, Office of Biological and Environmental Research of the U.S. Department of Energy, under contract DE-AC02-05CH11231. The other data collection was supported by Laboratory Directed Research and Development (LDRD) funding from Berkeley Lab, provided by the Director, Office of Science, of the U.S. Department of Energy under contract DE-AC02-05CH11231 and a Community Science Project from the Joint Genome Institute to M.J.B., J.B., A.P.A., and A.D. The work conducted by the U.S. Department of Energy Joint Genome Institute, a DOE Office of Science User Facility, is supported by the Office of Science of the U.S. Department of Energy under contract no. DE-AC02-05CH11231.

## Author contributions

AMD, APA, MNP, MJB, and JB conceived the project. AMD, APA, MJB, and JB supervised the project. AMD led the experimental work. AMD, KMW, RJW, RAM, MC, JR, JVK, JSL, YS, ZE, and HS collected data. RC isolated bacteria. MNP and AMD analyzed the fitness data. RAM, RJW, and MNP assembled genomes. BER provided resources and advice on *S. elongatus* experiments. MNP, MJB, and AMD wrote the paper.

## Supplementary information

### In separate excel file

Supplementary Table 1 – List of essential genes

Supplementary Table 2 – Growth on carbon substrates

Supplementary Table 3 – Growth on nitrogen substrates

Supplementary Table 4 – Growth on stress compounds

Supplementary Table 5 – Catabolic genes in *Cupriavidus basilensis*

Supplementary Table 6 – *E. coli* genes with specific phenotypes in carbon and nitrogen source experiments

Supplementary Table 7 – Genes with conserved specific phenotypes or conserved cofitness

Supplementary Table 8 – Genes with conserved specific phenotypes under cisplatin stress

Supplementary Table 9 – Genes with specific phenotypes during D-xylose utilization Supplementary Table 10 – Reannotation of ABC transporters

Supplementary Table 11 – Uncharacterized protein families with conserved specific phenotypes or conserved cofitness

Supplementary Table 12 – Bacteria used in this study

Supplementary Table 13 – Media formulations used in this study

Supplementary Table 14 – Genome sequencing statistics

Supplementary Table 15 – Transposon mutagenesis details

Supplementary Table 16 – Genome annotation summary

**Supplementary Figure 1:**
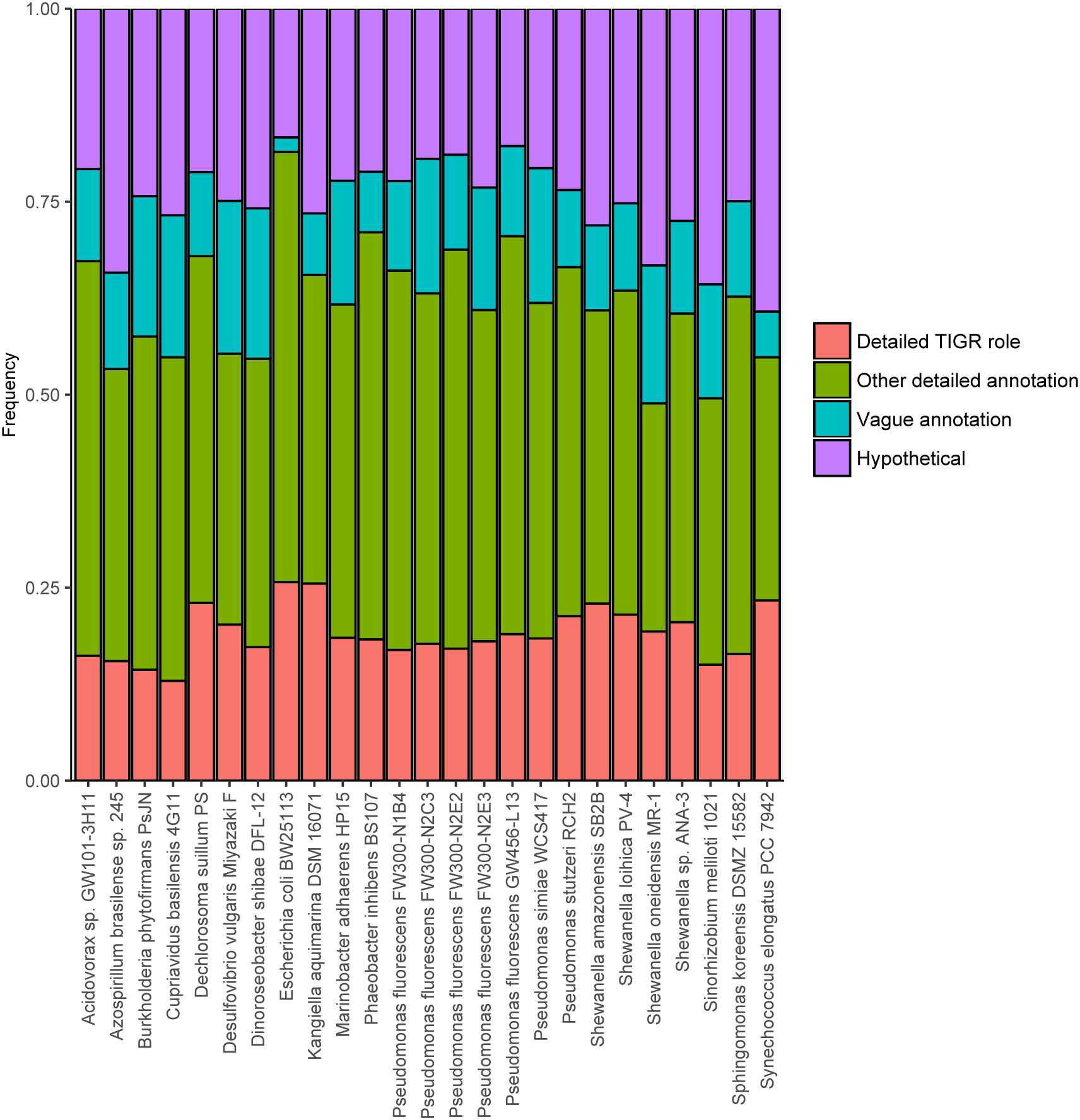
Protein annotations for the 25 bacteria. The fraction of proteins in each genome with different levels of annotation (see main text and Methods) is color-coded.

**Supplementary Figure 2:**
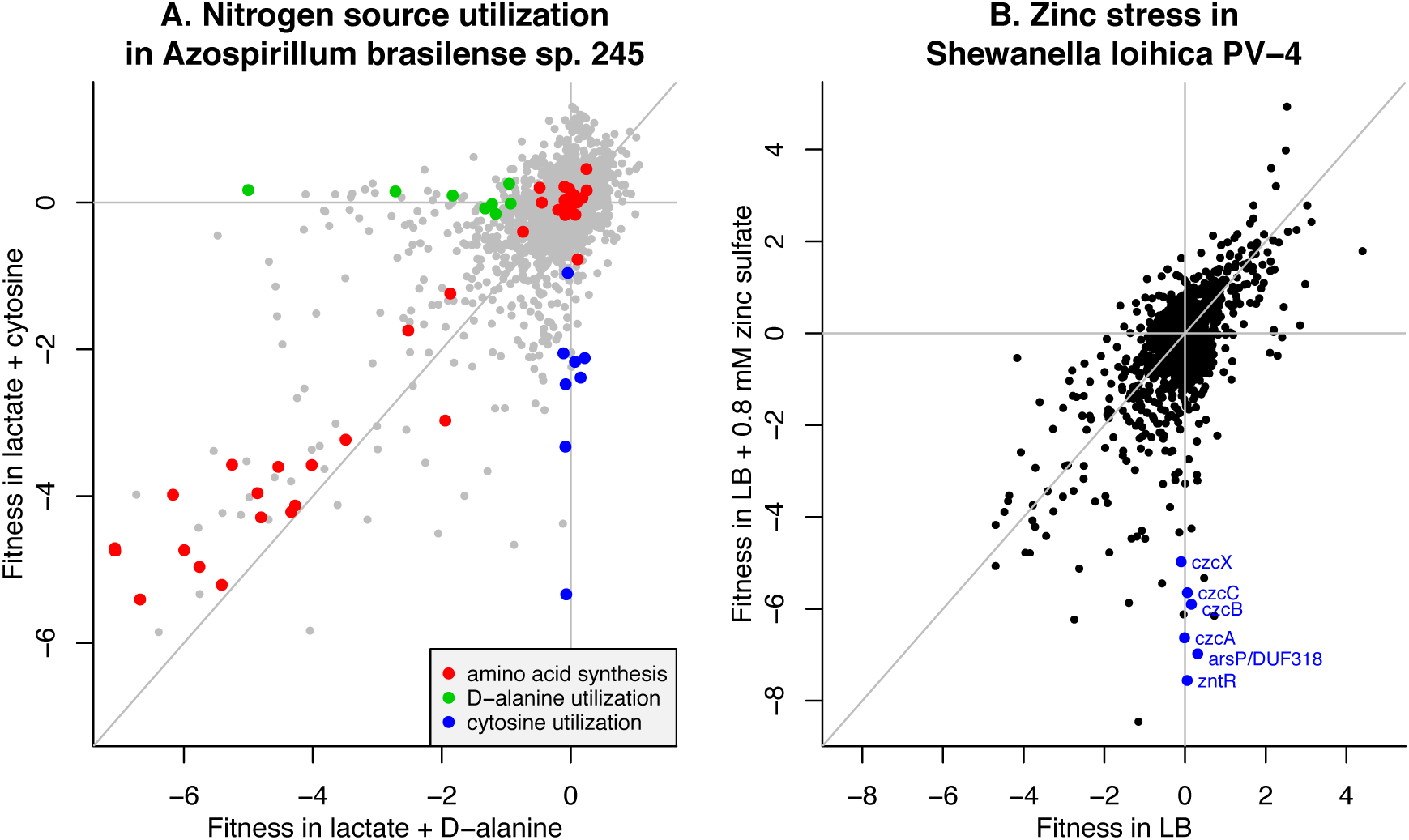
Examples of nitrogen source and stress fitness experiments. (A) The utilization of D-alanine or cytosine by *Azospirillum brasilense* sp. 245. Each point shows the fitness of a gene in the two conditions. The data is the average of two biological replicates for each nitrogen source. Amino acid synthesis genes were identified using the top-level role in TIGRFAMs. The genes for D-alanine utilization were a D-amino acid dehydrogenase (AZOBR_RS08020), an ABC transporter operon (AZOBR_RS08235:RS08260), and a LysR family regulator (AZOBR_RS21915). The genes for cytosine utilization were cytosine deaminase (AZOBR_RS31895) and two ABC transporter operons (AZOBR_RS06950:RS06965 and AZOBR_RS31875:RS31885). (B) Zinc stress in *Shewanella loihica* PV-4. We compare fitness in rich media with added zinc (II) sulfate to fitness in plain rich media. The LB data is the average of two biological replicates. The highlighted genes include a putative heavy metal efflux pump (CzcCBA or Shew_3358:Shew_3356), a hypothetical protein at the beginning of the *czc* operon (CzcX), a zinc-responsive regulator (ZntR or Shew_3411), and another heavy metal efflux gene related to *arsP* or DUF318 (Shew_3410). *CzcX* lacks homology to any characterized protein, but homologs in other strains of *Shewanella* are also specifically important for resisting zinc stress. In both panels, the lines show x = 0, y = 0, and x = y.

**Supplementary Figure 3.**
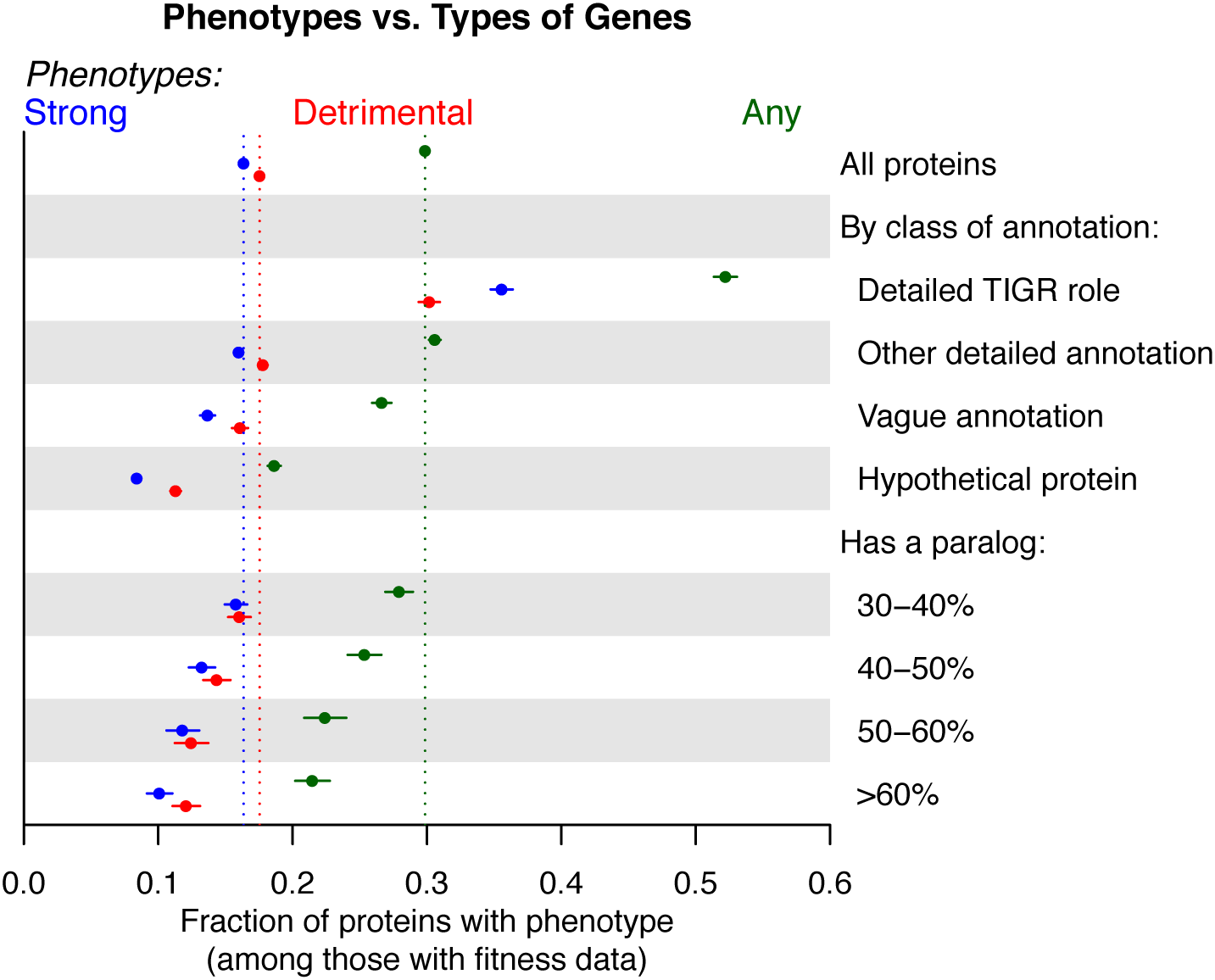
Phenotypes versus types of genes. We categorized proteins in our data set by their type of annotation or by whether they have homologs in the same genome (“’paralogs”). For each category, we show the fraction of genes that have significant phenotypes, and more specifically the fractions that have strong phenotypes (fitness < −2 and *t* < −5) or are detrimental to fitness (fitness > 0).

**Supplementary Figure 4.**
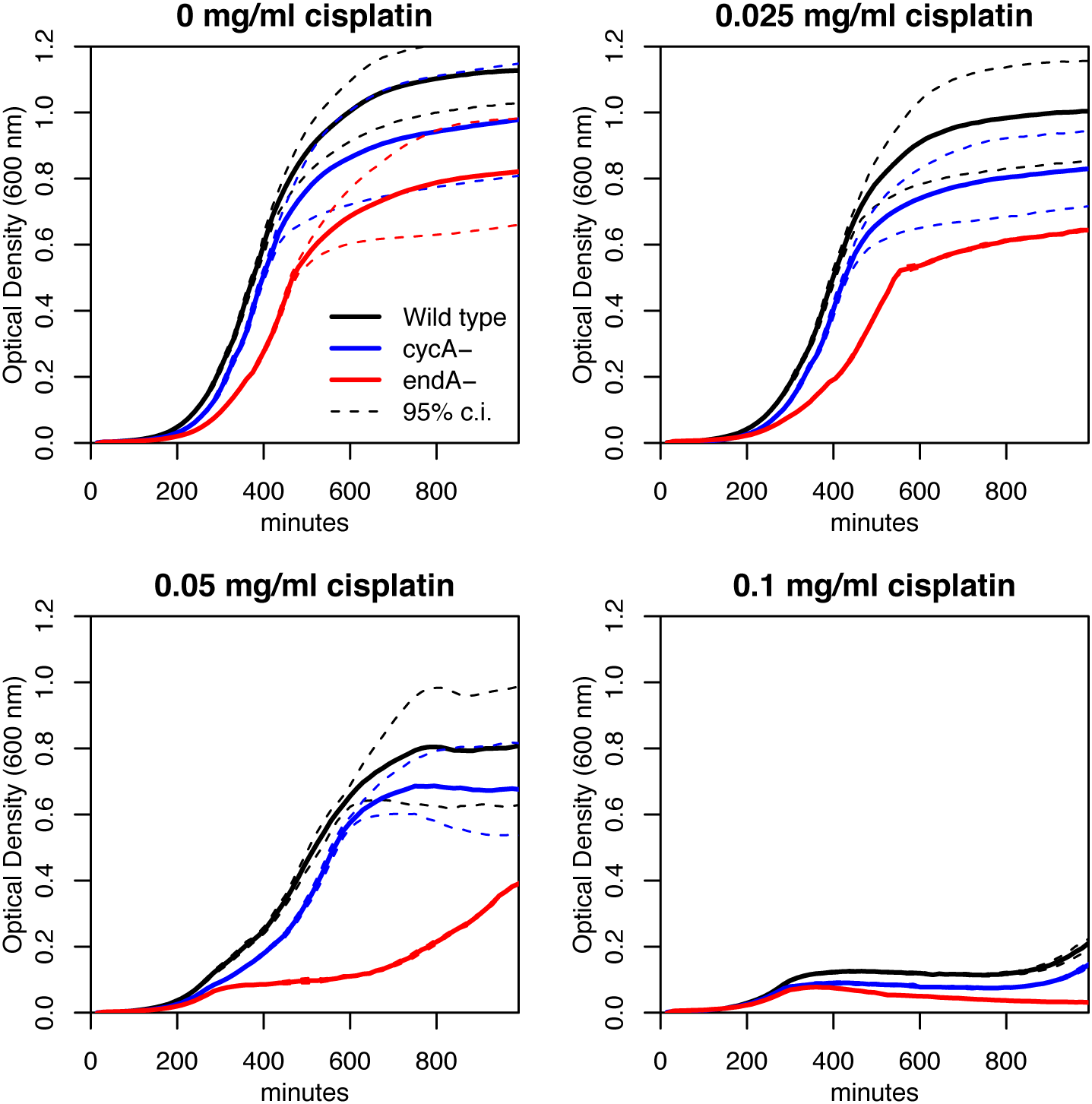
EndA is important for cisplatin resistance. We compared the growth of the *endA-* deletion strain from the Keio collection to the growth of the base strain (BW25113) and the growth of a *cycA-* deletion strain (*cycA* encodes an amino acid transporter and is not expected to have a phenotype in this condition.) All experiments were conducted at 30°C in LB media with varying levels of cisplatin added. This experiment was conducted in a 96-well plate. Each growth curve is the average of 12 replicate wells and the dashed lines show 95% confidence intervals. *endA* encodes endonuclease I, which is believed to be in the periplasm -- it is released from cells after various shocks ^1,2^. There are also mutants of *E. coli* that leak several periplasmic enzymes including some endonuclease I ^3^. EndA has a likely signal peptide and no other transmembrane helix, which is consistent with being in the periplasm. It remains unclear how EndA plays a role in the response to DNA damage if it is located in the periplasm.

**Supplementary Figure 5.**
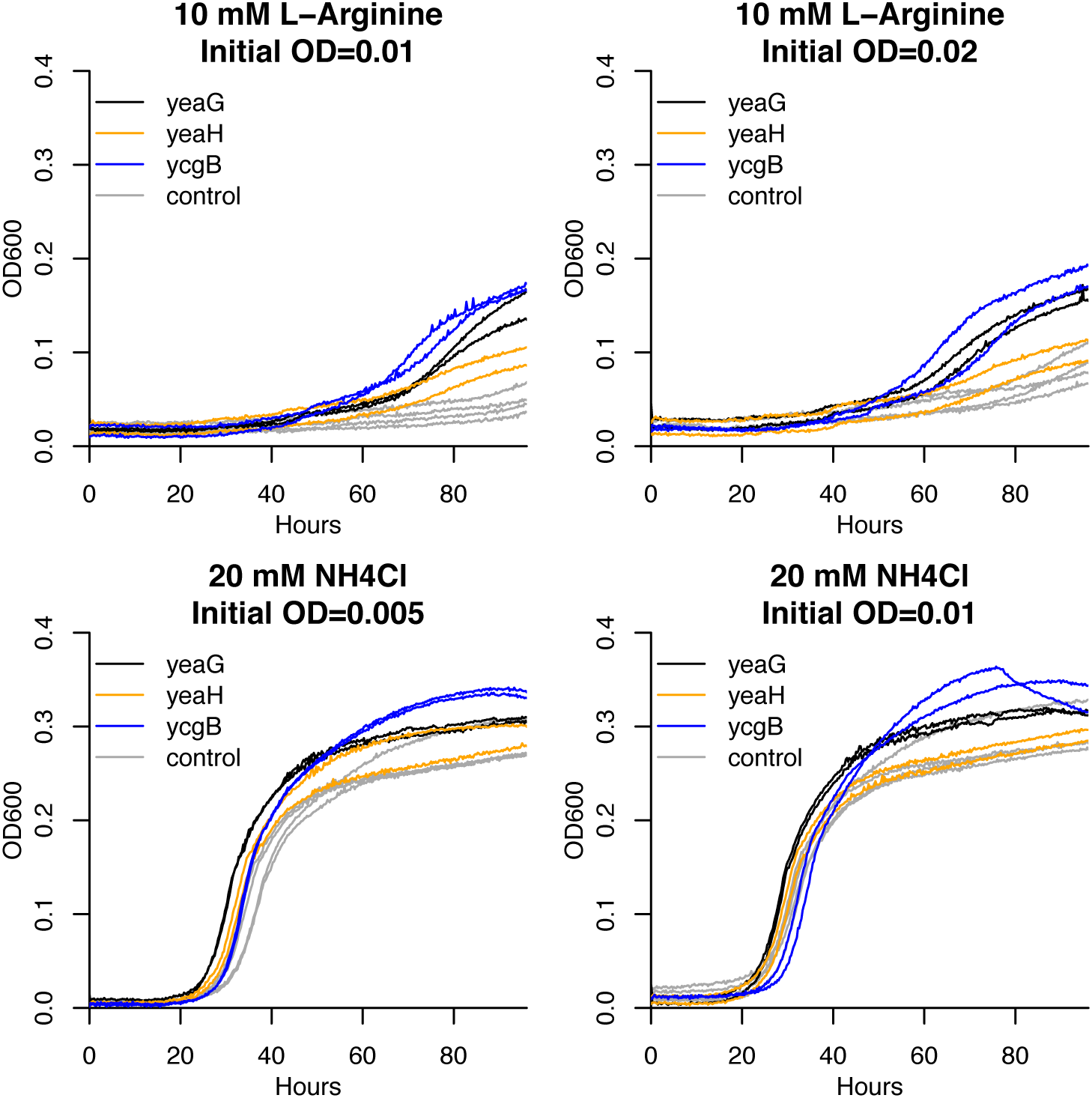
Growth of signaling mutants in *Escherichia coli*. We grew deletion strains from the Keio collection ^4^ in M9 glucose media with varying nitrogen sources. The signaling mutants are in *yeaG*, *yeaH*, and *ycgB*. Control mutants are deletions of two pseudogenes, *agaA* or *ygaY*. The signaling mutants had a strong growth advantage on L-arginine, as expected from fitness assays with 10 mM L-arginine as the nitrogen source.

**Supplementary Figure 6.**
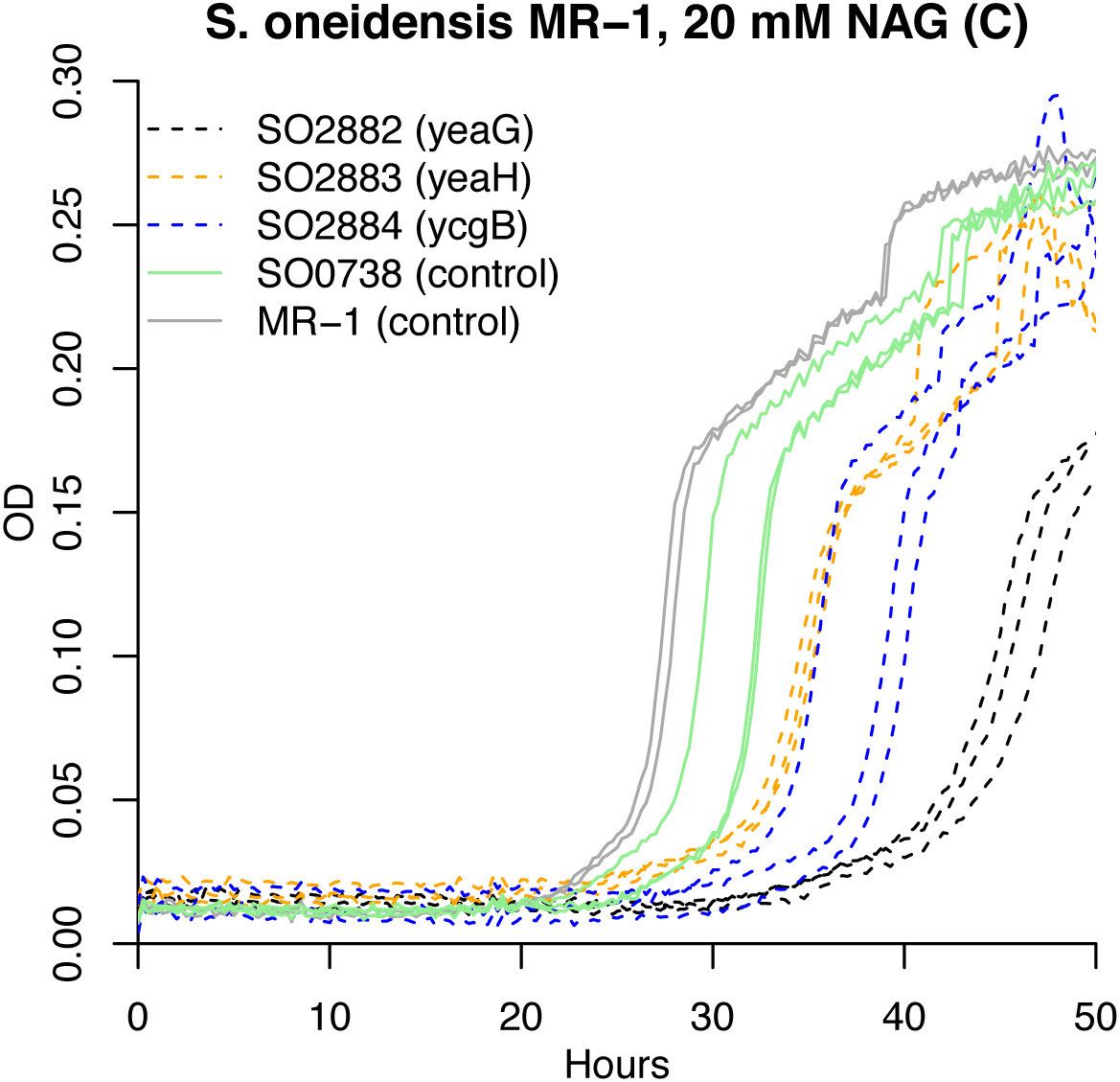
Growth of signaling mutants from *Shewanella oneidensis*. We grew various strains derived from *S. oneidensis* MR-1 in a defined medium (ShewMM_noCarbon) with 20 mM N-acetylglucosamine (NAG) as the carbon source. This medium contains ammonium as the nitrogen source. These mutants have transposons inserted within the signaling pathway (SO2882:SO2884) or in a pseudogene (SO0738, identified as such by Romine and colleagues ^5^). We also grew wild-type *S. oneidensis* MR-1 as an additional control. Insertions in the signaling pathway had a increased lag and a slower growth rate, as expected from fitness assays in this condition. The mutants are from a previously described collection ^6^ and were generated independently from the mutants that were used to generate the fitness data.

### Supplementary Note 1. Validation of essential genes analysis

Our approach to identify essential genes was initially validated for *Synechococcus elongatus* ^7^. To verify that this approach gave reasonable results for other bacteria, we compared our list of putatively essential proteins in *E. coli* K-12 to genes that are essential ^8^ or important for growth in LB ^4^. We also compared the list of essential proteins in *Shewanella oneidensis* MR-1 to a previous analysis based on a different set of transposon mutants ^6^. Finally, for all of the genomes, we examined the functional categories according to TIGRFAMs ^9^ and whether essentiality was conserved.

Of the 259 essential proteins in *E. coli*, 219 (85%) were in our list of essential proteins. Our list also included another 122 proteins (from 3,887 non-essential proteins), which corresponds to a false-discovery rate of 36%. Of those 122, 25 have reduced growth in LB, with OD_600_ after 22 hours being in the bottom 10% of genes ^4^. If we consider these as important genes then the FDR drops to 29%. The remaining false positives tend to be somewhat shorter than true essential genes, with median lengths of 638 and 942 nucleotides, respectively. These 97 “false positives” include 10 genes that are involved in protein synthesis or protein fate as well as other genes that are likely to be important for fitness (*minE*, *mviN*, *dapF*, *holD*, *ubiB*, *ubiD*, and *purB*).

In *S. oneidensis* MR-1, our list contained 397 proteins. We compared this to a list of essential genes that we had previously generated using a different transposon and Sanger sequencing ^6^. The previous analysis was conservative as genes that were not expected to be essential (based on orthology to *E. coli* or *Acinetobacter*, which are also γ-Proteobacteria) were required to be adjacent to another essential gene or to be conserved across most other *Shewanella* genomes. Of the 397 proteins in our new list, 298 were previously classified as essential and 83 were classified as unknown, Just 16 were previously identified as dispensable, and 2 of these are genes that are essential in *E. coli*. This implies a false discovery rate of around 16/(397-83) = 5%, with the caveat that insertions in a gene might be selected against even though the gene is not itself important (i.e., polar effects). We also examined the 15 proteins that are putatively essential in *S. oneidensis* MR-1 but lacked clear orthologs in the closely related strain *S. sp.* ANA-3, as these might be more likely to be false positives. Of these 15, at least seven are plausibly essential, including three prophage repressors that are probably required to prevent prophage excision; a gene adjacent to one of these repressors; the RepA protein that is probably required to maintain the megaplasmid, a tRNA synthetase (SO3128.2) with a putative internal stop codon that nevertheless forms full-length protein ^10^, and ribosomal protein L25 (whose ortholog was missed because it seems to be annotated with the wrong start codon).

We then considered TIGR roles that are likely to be essential. The roles we chose were: DNA metabolism; transcription; protein synthesis; protein fate; energy metabolism; cell envelope; fatty acid and phospholipid metabolism; purines, pyrimidines, nucleosides, and nucleotides; amino acid biosynthesis; and biosynthesis of cofactors, prosthetic groups, and carriers. We included biosynthetic and energetic genes as likely-essential roles because many bacteria cannot take up as wide a range of nutrients as *E. coli* can or have more limited ways of creating energy. In *E. coli* and *S. oneidensis*, these categories account for 62-67% of putatively essential genes but just 12-18% of other genes. In the 25 bacteria, these categories accounted for 44-69% of putatively essential genes but just 8-17% of other genes. This confirms that in each bacterium, most of these genes are essential.

Finally, we asked whether the putatively essential proteins had orthologs in other bacteria that were essential. Overall, 84% of the essential proteins were confirmed by conservation (they had an ortholog that was also essential). The organisms with the lowest proportions of conserved essentials were *S. elongatus* and *D. vulgaris*(56% and 64%, respectively). This might reflect their unique sources of energy (photosynthesis and dissimilatory sulfate reduction, respectively) or their evolutionary distance from the other bacteria that we studied.

### Supplementary Note 2. Rationales for annotating domains of unknown function

#### Specific Predictions

##### DUF485 (PF04341): component of actP-like carboxylate transporters

Several representatives of DUF485 (which is known as *yjcH* in *E. coli*) are cofit with an adjacent ActP-like permease. Examination of the per-strain data did not show any evidence of polar effects, so we suggest that DUF485 is required for the activity of the permease. The paper that characterized ActP in *E. coli* did not rule out the requirement of another gene ^11^. DUF485 seems to be a membrane protein (i.e., AZOBR_RS02935 has two transmembrane helices and a possible signal sequence). Together, this suggests that DUF485 is a component of the transporter. The clearest phenotypes are for pyruvate utilization (i.e. AZOBR_RS02935).

##### DUF1513 (PF07433): outer membrane component of ferrous iron uptake

DUF1513 is in a conserved operon and is cofit with the other genes in that operon in multiple bacteria. The operon contains EfeO-like, DUF1111, a second EfeO-like, and DUF1513. EfeO has an unknown role in iron uptake by *efeUOB* and is a putative periplasmic lipoprotein. DUF1111 is proposed to be a homolog of EfeB, a di-heme peroxidase involved in ferrous iron uptake ^12^. Thus, the operon seems to be involved in ferrous iron uptake. Related operons in α-Proteobacteria often contain bacterioferritin, which is consistent with that role.

DUF1513 has a putative signal peptide (PSORTb) and has similarity to beta propellers with 6-8 repeats (Pfam clans). This suggests that it is the outer membrane component of this system.

Although the fitness data shows that DUF1513 is involved in this process, it is not certain that ferrous iron is the substrate. Several representatives of DUF1513 are pleiotropic, but AO356_18450 is specifically important for chlorite resistance and Psest_1156 is sensitive to various metals. Inhibiting iron uptake would plausibly create these phenotypes.

##### DUF1656 (PF07869): component of efflux pump with MFP and FUSC

Several representatives of DUF1656 are in an operon with and cofit with a RND efflux pump and a fusaric acid resistance-like protein. DUF1656 is related to *E. coli* YdhI and AaeX/YhcR. Homologs of these proteins are sometimes annotated as inner membrane efflux pump components or as Na+-dependent SNF-like transporters but we could not find the rationale for these annotations. (As of March 2016, EcoCyc reports that the functions of *aaeX* and *ydhI* are not known.)

This family has a variety of stress sensitivity phenotypes. In *Cupriavidus basilensis*, this efflux pump is specifically important for the utilization of 4-hydroxybenzoate. It could be involved in uptake, or 20 mM of 4-hydroxybenzoate may have been inhibitory and it is involved in efflux. Similarly, a number of representatives are important for the utilization of octanoate and it is not clear if this reflects uptake or efflux. In several strains of *Pseudomonas fluorescens*, the efflux pump is either important for or detrimental during acetate utilization. Finally, in *Zymomonas mobilis*, we found that a similar system (ZMO1432-ZMO1431-ZMO1430) is involved in resisting hydrolysate ^13^.

##### DUF1854 (PF08909): subunit of transporter for efflux of an amino acid polymer

In two β-Proteobacteria, *Acidovorax* sp. 3H11 and *Cupriavidus basilensis* 4G11, representatives of DUF1854 (i.e., RR42_RS04420) are cofit with two nearby genes that are annotated as cyanophycin synthetases as well as an ABC-like transporter. Cyanophycin is a copolymer of aspartate and arginine that many cyanobacteria use to store nitrogen; it is also formed by some heterotrophic bacteria ^14^. The putative cyanophycin synthetases (ChpA and ChpA') have been studied in another strain of *Cupriavidus necator* and do not form cyanophycin but may produce a different light-scattering polymer ^15^. Also, the genomes of many β-Proteobacteria contain *chpA* and *chpA'* but do not appear to encode the cyanophycinase (*chpB*) to break down the polymer, which hints at a different role for these genes ^14^. As *chpA* and *chpA'* were cofit with an ABC transporter in both organisms, we propose that the polymer is being exported to form part of the cell wall rather than serving as a storage compound. This can also explain why the genes are important for motility (in *Acidovorax*) or have pleotropic stress phenotypes (in both organisms). (We did not succeed in measuring fitness during motility in *C. basilensis*.) The ABC transporter contains an ABC transporter transmembrane domain and an ATPase domain. DUF1854 is usually found in an apparent operon with the ABC transporter, but in a few genomes the two proteins are fused together, as pointed out by the Pfam curators. Together with the cofitness, this suggests that DUF1854 is involved in the transport and forms a complex with the ABC transporter.

##### DUF2849 (PF11011): electron source for sulfite reductase

This family is usually upstream of *cysI*, the beta subunit of sulfite reductase. These sulfite reductases are important for fitness in our defined media, which confirms that they are indeed sulfite reductase and not nitrite reductase, as sulfate is the sulfur source in these media and sulfate must be reduced to sulfite and then to sulfide before it is assimilated. (In *S. meliloti*, *cysI* or SMc02124 was misannotated as “nitrite reductase.”) However these genomes do not seem to contain the alpha flavoprotein subunit (*cysJ*). Instead, DUF2849 is found upstream. Several representatives of DUF2849 have similar fitness patterns as the downstream *cysI*(SMc01054, AZOBR_RS10130, Ga0059261_1497). The phenotypes of DUF2849 do not seem to be due to polar effects, as strains with insertions in either orientation have low fitness in defined media. Also, DUF2849 is found fused to *cysI* in *Pseudomonas putida* GB-1. Altogether this suggests that DUF2849 is required for the activity of sulfite reductase. Usually *cysJ* is the electron source for *cysI* so we propose that in its absence, DUF2849 fulfills this role. (The relationship between the representatives of DUF2849 with similar cofitness was missed by the automated analysis of orthologs, as these alignments just missed the cutoffs of coverage > 80% or E < 10^−5^. So this case is absent from **Supplementary table 11**.)

##### DUF4212 (PF13937, TIGR03647): small subunit of transporter for D-alanine, lactate

DUF4212 was predicted by the TIGRFAM curators to be the small subunit of a solute:sodium symporter because of its hydrophobicity and conserved gene context. Multiple members of this family (e.g., Sama_1522 or Psest_0346) are cofit with a putative large subunit of a symporter that is downstream (e.g., Sama_1523 or Psest_0347). Sama_1522 from *Shewanella amazonensis* SB2B is important for fitness when D,L-lactate is the carbon source; similarly, a homolog in *S. oneidensis* MR-1 (SO2858) is more mildly important in some D,L-lactate conditions. Psest_0346 in *Pseudomonas stutzeri* RCH2 (51% identical to Sama_1523) is very important for D-alanine utilization and strongly detrimental during L-alanine utilization. Note that alanine and lactate are chemically analogous three-carbon organic acids, with alanine having an amino group where lactate has a hydroxyl group. Overall, our data confirms that that DUF4212 is required for the transporter's activity, so it is probably an additional subunit. Some members of DUF4212 are cofit with a nearby member of COG2905 (RNAse T domain protein) (e.g., Sama_1525 or Psest_0349); we speculate that COG2905 is required for expression of the symporter.

##### UPF0060 (PF02694, COG1742): efflux pump for thallium (I) ions

UPF0060 is specifically important for resisting thallium (I) stress in *Cupriavidus basilensis* (RR42_RS34240), in *Sphingomonas koreensis* (Ga0059261_1942), and in two strains of *Pseudomonas fluorescens*. This family includes E. coli YnfA, which is an integral membrane protein. Nir Hus (PhD dissertation, 2005) proposed that YnfA belongs to the SMR (small multi-drug-resistance) family and SCOOP ^16^ (a family-family relationship finder) shows similarity to transporters and efflux pumps. UPF0060 is sometimes adjacent to cation efflux genes related to *czcD* or *zntA*. So, we propose that these members of UPF0060 are efflux pumps for thallium (I).

##### UPF0126 (PF03458, COG2860): glycine transporter

A number of representatives of UPF0126 are specifically important for utilizing glycine (PGA1_c00920, SO1319, Sama_2463, Psest_1636, AO353_13110). These genes contain a pair of UPF0126 domains and are predicted to be membrane proteins. SCOOP identified similarity between UPF0126 and TRIC channels, so we propose that this is a family of glycine transporters. This family includes *yicG* from *Escherichia coli*, which is not characterized (and which we did not identify any significant phenotypes for).

#### Pathway-level Predictions

##### DUF444 (yeaH, PF04285) and SpoVR (ycgB, PF04293): signaling with serine kinase yeaG

As discussed in the main text, we identified strong and conserved cofitness between *yeaG*, *yeaH*, and *ycgB* in many different bacteria, including *Escherichia coli*. These genes form a single operon in most bacteria, but in *E. coli* they are broken up into two operons (*yeaGH* and *ycgB*). The biological role of YeaG is not known but it is a PrkA-like protein kinase and can phosphorylate itself or casein *in vitro* ^17^. SpoVR is so named because a member of this family is involved in sporulation in *Bacillus subtilis,* but nothing is known about its biochemical function and the organisms that we studied do not sporulate. Since YeaG appears to be a signaling protein, we infer that YeaH and YcgB are involved in this signaling pathway as well. The mutant phenotypes of these genes are not conserved, but there is a potential commonality relating to the utilization of amino acids: these genes are detrimental for the utilization of L-arginine as the nitrogen source by *E. coli*, detrimental for the utilization of L-methionine or L-phenylalanine as the nitrogen source by *S. oneidensis* MR-1, and detrimental for the utilization of L-serine as the carbon source by *P. fluorescens* FW300-N2E3. So we suspect that the ultimate target(s) of this signaling pathway is involved in nitrogen metabolism.

##### DUF466 (PF04328, COG2879): accessory to pyruvate transport by cstA-like

Three representatives of DUF466 were cofit with a nearby cstA-like protein. In *E. coli*, CstA is reported to be a peptide transporter. In the *Desulfovibrio alaskensis* G20, a cstA-like protein (Dde_2007) is specifically important for fitness when pyruvate is the carbon source (data from ^18,19^). The only strong phenotype for the DUF466 and *cstA* in our data were in *C. basilensis*, where they were specifically important for pyruvate utilization. So, we propose that DUF466 is required for pyruvate transport by a CstA-like protein.

##### DUF692 (PF05114), DUF2063 (PF09836): chlorite stress signaling proteins

DUF692 and DUF2063 form a conserved operon that is important for chlorite resistance in *P. stutzeri* RCH2 (Psest_0116:Psest_0117), *Kangiella aquimarina* (B158DRAFT_1333:B158DRAFT_1334), and *Shewanella amazonensis* SB2B (Sama_1305 = DUF692; DUF2063 seems to have been replaced by a hypothetical protein, Sama_1304, which also has this phenotype). In *S. amazonensis*, DUF692 is cofit with the hypochlorite scavenging system *yedYZ* (Sama_1893:Sama_1892, see ^20,21^), so DUF692 probably has another role rather than detoxifying hypochlorite directly. DUF692 is a putative xylose isomerase or epimerase-like and DUF2063 is a putative DNA binding protein. We propose that these genes are involved in sensing an aspect of chlorite stress or metal stress. Some homologs are in an operon with an extracellular type sigma factor that responds to heavy metal stress (e.g. NGO1944 from *Neisseria gonorrhoeae*, which regulates methionine sulfoxide reductase). This suggests that DUF2063 might be an anti-anti-sigma factor rather than a DNA-binding protein. Consistent with this, in *Neisseria*, there is a putative anti-sigma factor downstream of the sigma factor (e.g., NMB2145), which is not similar to DUF2692 or DUF2063.

##### DUF934 (PF06073): accessory protein for sulfite reduction

This uncharacterized protein family is conserved downstream of *cysI*, the β subunit of sulfite reductase. In SEED, members of this family are annotated as CysX or “Oxidoreductase probably involved in sulfite reduction,” but we were not able to find any published experimental evidence. They were probably annotated based on conserved gene proximity. (DUF934 does not seem to be homologous to the small ferredoxin-like CysX of *Corynebacterium glutamicum* described by Ruckert et al ^22^; furthermore, it lacks the CxxC motifs that are conserved in CysX.) We found that DUF934 was strongly cofit with various genes in the sulfate assimilation pathway in various *Pseudomonas* (i.e., Psest_2088) as well as in *Marinobacter adhaerens* HP15 and *C. basilensis* 4G11. In *M. adhaerens* and in *C. basilensis*, there are other sulfate assimilation genes downstream, but not in the *Pseudomonas* species, so this is not a polar effect.

##### DUF971 (PF06155): FeS cluster maintenance protein

In several *Shewanella* or *Pseudomonas* species, a representative of DUF971 is cofit with Mrp, BolA, and/or YggX. All of these proteins are related to FeS cluster maintenance: Mrp is an FeS loading protein; BolA is related to the Fra2 protein of *Saccharomyces cerevisiae* which is an FeS cluster protein and regulates iron levels; and YggX plays a role in oxidation resistance of FeS clusters. DUF971 is pleiotropic but a number of representatives are important for resisting cobalt (II) or paraquat; either of these might disrupt FeS clusters. Nothing is known about the biochemical function of DUF971, but in eukaryotes, DUF971 is found as a domain within gamma-butyrobetaine hydroxylase and trimethyllysine dioxygenase proteins, and it is also found within the chloroplast 4Fe4S cluster scaffold protein HCF101 (fused with an Mrp-like domain). These occurrences are consistent with DUF971 having a role in FeS cluster maintenance.

##### DUF1302 (PF06980), DUF1329 (PF07044): export of cell wall component

We identified cofitness for a gene cluster comprising DUF1302 (e.g., Sama_1588 in *Shewanella amazonensis* SB2B), DUF1329 (Sama_1589), a BNR repeat protein (COG4447; Sama_1590), and a putative RND export protein (COG1033; Sama_1591). The BNR repeat protein is related to a photosystem II stability/assembly factor (ycf48 or hcf136) and has a beta-propeller fold (PDB 2xbg). These genes were conserved near each other and are cofit in *Kangiella aquimarina* DSM 16071 as well. In *Pseudomonas stutzeri* RCH2, the cluster is broken up into two pieces, Psest_1122:_1123 (DUF1302 and DUF1329) and Psest_1923:_1924 (BNR repeat protein and RND export protein).

Again these proteins are cofit, although in one condition (tyrosine as a carbon source), the two groups have strong and opposing phenotypes; this could imply some separation of function, or it could relate to the existence of paralogs for DUF1329 in this organism. These genes were cofit in some strains of *Pseudomonas fluorescens* as well. In both *S. amazonensis* and *P. stutzeri*, their phenotypes are pleotropic.

Several lines of evidence suggest that these proteins are in the outer membrane and affect the cell wall. DUF1329 has some similarity to the outer membrane protein LolB (e.g., the C-terminal part of Sama_1589 is similar to CATH superfamily 2.50.20.10). In *Delftia sp.* Cs1-4, both DUF1302 and DUF1329 are reported to be components of extended outer membrane vesicles or nanopods ^23^. BNR repeat proteins are often found in the outer membrane. And a mutant of a member of DUF1329 in *P. fluorescens* F113 is reported to have increased swimming motility ^24^. We propose that these four proteins work together to export a component of the cell wall.

Incidentally, a number of other studies have discussed the DUF1329 family but without identifying a biochemical function or a mutant phenotype. In *Delftia sp.* Cs1-4, a DUF1329 family member (phnK) is in a genomic island for phenanthrene catabolism, where it is also near a RND efflux protein ^25^. In *Thauera aromatica* T1, expression of a member of this family (*pipB*) is induced by p-cresol (M. Chatterjee, PhD Thesis 2012). (A DUF1302 member, *pipA*, was also induced.) In *Comamonas testosteroni*, ORF61 (DUF1329) is in a gene cluster for steroid degradation but is not required for it ^26^. The presence of DUF1329 in these gene clusters seems to suggest a specific role in the maturation of an outer membrane transporter, or directly in the transport of aromatic compounds, but this is not what we observed in *S. amazonensis* or *P. stutzeri*.

##### DUF1654 (PF07867): DNA repair protein

In several strains of *Pseudomonas fluorescens*, members of DUF1654 (PF07867) are specifically important for resisting the DNA-damaging agent cisplatin (e.g., AO356_00980). Furthermore, these genes are upstream of an endonuclease precursor, and close homologs of these proteins are predicted to be regulated by LexA, which controls the DNA damage response (e.g., PFL_2098 from *P. fluorescens* Pf-5). These imply a role for DUF1654 in DNA repair, but we have little idea of its molecular function. Polar effects seem unlikely because DUF1654 has a strong phenotype relative to the downstream endonuclease, but we cannot rule them out entirely. DUF1654 has some similarity (SCOOP) to an anti-parallel beta barrel family (calycin_like, PF13944).

##### DUF2946 (PF11162): copper homeostasis protein

In two strains of *Pseudomonas fluorescens*, DUF2946 is in an operon with and cofit with a TonB-dependent copper receptor (TIGR01778) and a Cox17-like copper chaperone (i.e., AO356_08325 with _08320 and _08330; the copper chaperone is also referred to as CopZ). Some members of DUF2946 are labeled as being homologous to COG2132, which includes *E. coli* CueO and SufI, but those are much longer proteins (around 500 amino acids, while DUF2946 is around 100 amino acids). We propose that DUF2946 is also involved in copper uptake or homeostasis.

##### DUF3584 (PF12128): smc-like chromosome partitioning protein

In three species of *Shewanella*, DUF3584 was specifically important for resisting cisplatin stress (i.e., Sama_0592) and also for motility. According to HHSearch, DUF3584 is related to the N-terminal domain of Smc, or RecF or RecN, which are all involved in DNA repair or recombination. Together, this strongly suggests that DUF3584 is involved in chromosome segregation or DNA repair, but we do not have a specific proposal for its biochemical role. DUF3584 is conserved cofit with two hypothetical proteins (e.g., Sama_0594 and Sama_0593) that do not belong to any annotated families, and the three genes probably act together.

##### UPF0227: hydrolase affecting the cell envelope

UPF0227 includes the *E. coli* protein YqiA. *In vitro*, YqiA has esterase activity on palmitoyl-CoA and p-nitrophenylbutyrate ^27^, but little is known about its biological role. Mutants in this family have diverse phenotypes, including sensitivity to bacitracin, which inhibits synthesis of a peptidoglycan precursor (*E. coli* yqiA, Sama_3044, and Psest_0482), and sensitivity to carbenicillin, which a β-lactam antibiotic and inhibits peptidoglycan synthesis (SO3900, Psest_0482). In some *Shewanella* species (but not in *S. oneidensis* MR-1), this family is important for fitness in many defined media conditions. These phenotypes are consistent with an effect on cell wall or membrane composition. Given its esterase activity on palmitoyl-CoA, an effect on lipid composition seems most likely. A caveat is that these genes are upstream of the essential protein *parE*, and it difficult to rule out a phenotype from reducing *parE* expression (a polar effect). Although we cannot rule out this explanation, we did find that insertions within *E. coli*'s *yqiA* or within Sama_3044 have strong phenotypes in either orientation, which makes it less likely.

